# High content 3D imaging method for quantitative characterization of organoid development and phenotype

**DOI:** 10.1101/2021.03.26.437121

**Authors:** Anne Beghin, Gianluca Grenci, Harini Rajendiran, Tom Delaire, Saburnisha Binte Mohamad Raffi, Damien Blanc, Richard de Mets, Hui Ting Ong, Vidhyalakshmi Acharya, Geetika Sahini, Victor Racine, Remi Galland, Jean-Baptiste Sibarita, Virgile Viasnoff

**Affiliations:** Mechanobiology Institute, National University of Singapore, 117411, Singapore; Biomedical Engineering Department, National University of Singapore, 117583, Singapore; Univ. Bordeaux, CNRS, Interdisciplinary Institute for Neuroscience, IINS, UMR 5297, F- 33000 Bordeaux, France; QuantaCell, 33600 Pessac, France; Department of Biological Sciences, National University of Singapore, 117583, Singapore; IRL 3639 CNRS, 117411 Singapore

**Author notes:** Corresponding authors: Anne BEGHIN, Virgile Viasnoff.

## Abstract

Quantitative analysis on a large number of organoids can provide meaningful information from the morphological variability observed in 3D organotypic cultures, called organoids. Yet, gathering statistics of growing organoids is currently limited by existing imaging methods and subsequent image analysis workflows that are either restricted to 2D, limited in resolution, or with a low throughput. Here, we present an automated high content imaging platform synergizing high density organoid cultures with 3D live light-sheet imaging. The platform is an add-on to a standard inverted microscope. We demonstrate our capacity to collect libraries of 3D images at a rate of 300 organoids per hour, enabling training of artificial intelligence-based algorithms to quantify the organoid morphogenetic organization at multiple scales with subcellular resolution. We validate our approach on different organotypic cell cultures (stem, primary, and cancer), and quantify the development of hundreds of neuroectoderm organoids (from human Embryonic Stem Cells) at cellular, multicellular and whole organoid scales.

## INTRODUCTION

In organotypic 3D cell cultures, referred hereafter as organoids, stem cells differentiate and self-organize into spatial structures that share strong morphological and functional similarities with real organs. Organoids offer valuable models to study human biology and development outside of the body^1,2,3^. A growing number of organoids are being developed that mimic liver, brain, kidney, lung, and many other organs ^2,4,5^. Organoid differentiation is directed by the addition of soluble growth factors and extracellular matrix in a precise sequence during organoid development. However, in marked contrast to organs the development of organoids is quite heterogeneous.

Beyond numerous biological challenges^6,7^, the organoid cultures also pose technological challenges, in terms of cell culture methods, characterization of transcriptomics, and imaging. *In vivo* organ development occurs in a biological environment that results in a highly stereotypical self-organization of cell arrangements. Any phenotypic alteration can be used as a proxy to diagnose a diseased state. In contrast, organoids develop *in vitro* in minimally controlled micro-environments compatible with cell culture conditions, resulting in large variability in the development path and shape formation for each individual organoid. A recent study^8^ demonstrated that quantitative imaging of organoid shape (phenotype descriptors) coupled to the assessment of a few genetic markers allow definition of phenotypic development landscapes. They characterised the different organization paths that organoids can spontaneously adopt, and unravelled biological programs orchestrating the spatial induction of different cell fates. However, to achieve this goal, they had to accomplish a methodological and technological ‘tour-de-force’ ^9^ using a custom-made high throughput compatible solution for organoid growth, and commercially available high content imaging techniques. Arguably, the ability to relate the diversity of genomic expression in organoids with their phenotypic behaviour is a major step towards unleashing the full potential of organotypic cultures. It thus begs for the development of dedicated high content imaging approaches allowing characterization of organoid features at sub-cellular, multi-cellular, and whole organoid scales in 3D ^10,11^.

Optical sectioning fluorescent microscopy techniques, such as confocal or multiphoton laser scanning, hardly fit the needs of both high throughput and low phototoxicity/photobleaching. Commercially available high content screening platforms (e.g. PerkinElmer’s Opera or Molecular Devices’ ImageXpress), are usually adapted from 2D cell cultures and suffer from speed, penetration depth, bleaching, and image analysis issues when applied to 3D cultures. Light-sheet fluorescence microscopy (LSFM) is arguably the most appropriate imaging technique to visualize fine cellular details at the whole tissue level, with maximal speed and minimal phototoxicity and photobleaching. LSFM has been favored for the study of live organoids, as it allows long-term monitoring in physiological conditions. However, LFSM traditionally involves several objectives, and despite its superior imaging performances, it suffers from low imaging throughput. Recent studies using dual objective SPIM have tried to engineer solutions to improve the throughput^12^ using a liquid handling robot and complex dedicated imaging hardware. However, this solution does not permit long term sterile culture (more than 24 hours), yet periodic medium or extracellular matrix exchange is mandatory for organoid development and differentiation. To overcome these important limits, several approaches have been developed to perform LSFM with a single objective. High speed, high content SCAPE^13^ was used to obtain ultra-fast images of *in vivo* processes in individual *C. elegans* and *Zebra fish*. Alleviating the multi-objective constraint, other approaches proved capable to image multiple samples. Epi-illumination SPIM (ssOPM) enabled parallel 3D volume acquisition of single cell down to the single molecule level in 96 well plates^14^. ssOPM was used to screen 200 μm organoid in 3D to detect glucose uptake. The random positioning of the spheroids in the 96 well plate limited the study to 42 live samples.

It follows that existing technological solutions for 3D high content live imaging of organoids still remain ill-adapted for collecting large libraries of organoid data to be used to train artificial intelligence-based algorithms to later quantitatively assess and classify their morpho-dynamic profiles at the cellular, multicellular and whole organoid scale together.

In this context, we developed a versatile high-content screening (HCS) platform allowing streamlined organoid culture (from isolated hESC, hIPSC or primary cells to 3D multicellular differentiated organoids) and fast non-invasive 3D imaging. It integrates a next generation miniaturized 3D cell culture device, called the JeWells chip, that contains thousands of well-arrayed microwells (Jewell) flanked with 45° mirrors that allow fast, 3D high-resolution imaging by single objective light-sheet microscopy^15^. Compatible with any standard commercial inverted microscope after addition of a simple beam steering unit, our system enables imaging of 300 organoids in 3D with subcellular resolution and minimum photobleaching in less than one hour. As an illustration, we performed proof of concept experiments demonstrating that: 1) the JeWells chip allows long term culture (months) and differentiation protocols while precisely defining the number of initial cells in the wells. 2) Individual development of a large number of organoids can be monitored live using standard bright-field and 3D light-sheet fluorescence microscopy, in a HCS fashion. 3) Organoids can then be retrieved to perform further biological investigations, for example transcriptomic analysis. 4) The large number of acquired images can be used to train convolutional neural networks in order to precisely detect and quantify subcellular and multicellular features, such as mitotic and apoptotic events, multicellular structures (rosettes), and classify whole organoid morphologies.

## RESULTS

### Architecture of the disposable cell culture chips with integrated optics: JeWells

In order to streamline organoid culture and 3D high content imaging, we devised the JeWells chip, a new cell culture vessel optimized for 3D cell culture and light-sheet imaging. It allows the reproducible development and imaging of organoids in arrayed micro-cavities, with a good control of the initial cell number. The JeWells micro-cavities are moulded in a photocurable resin (NOA73) and shaped into truncated pyramids with 45° gold coated faces allowing the reflection of a laser beam to create a thin (2.5 µm) light-sheet perpendicular to the optical axis of a unique objective **(Figure 1a-b)**. The detailed protocol (Online Methods) for their microfabrication is illustrated in the **Supplementary Figure 1**. The Jewells are optimally arrayed in a glass bottom Petri dish, or a multi-well plate for fast screening. Their surface density is *150* Jewells/15 mm^2^, corresponding to the equivalent of 96 organoids in a single well of a 384 well-plate **(Figure 1-a)**. JeWells chips are passivated with Lipidure® that constitutes an enduring antifouling treatment over months (detailed protocols provided in Online Methods). The array of Jewells allows the simple and automatic identification of all individual organoids, facilitating their high content screening. The truncated pyramid shape provides four main advantages: *i-*it eliminates material loss during medium exchange by trapping the organoids (the pear in a bottle effect); *ii-*it provides a clear optical path for bright-field and high resolution fluorescent microscopy; *iii-*its 45° reflective surfaces allows multi-view single objective light-sheet fluorescence microscopy (soSPIM) on a conventional microscope; *iv*-it provides ideal beacons to automatically align the light-sheet and correct for drift. The Online Methods details of the acquisition set-up (**Supplementary Figure 1-2**). In brief, the 45° micromirrors of the Jewells embedding each organoid allow reflection of a laser beam **(Figure 1.b)**. Scanning the laser beam along a mirror (y-direction) creates a light-sheet perpendicular to the optical axis of the objective and illuminates a thin section of the sample. The fluorescent signal is then collected through the same unique objective. Translating the laser beam perpendicularly to the mirror (x-direction) allows translating of the light-sheet along the optical axis of the objective (z-direction). Synchronizing light-sheet position with the focal plane of the objective allows 3D optical sectioning. The integration of the optical components inside the chips eliminates the steric and alignment constraints of traditional multi-objective light-sheet microscopes. We devised an automatic acquisition pipeline that integrates an optional pre-screening step, the automated identification of the JeWells containing organoids to image, the precise positioning of the light-sheet and of the sample allowing, if needed, a drift correction during the acquisition (**Online Methods** and **Supplementary Figure 2**). It allows the rapid setup of the sequential 3D imaging of hundreds of organoids in one chip. We detailed the calibrations, the acquisition parameters, the light-sheet characteristics and the comparison with spinning disk in the Online Methods and in **Supplementary Figures 2-3**. In this following study, all results have been obtained with a 60x 1.27 NA magnification objective and opened pyramids of 70×70 µm upper aperture, 290×290 µm bottom-base and 110 µm height. We also validated the approach with 40x and 20x magnification objectives as a demonstration that our versatile technique allows changing of magnification depending on the biological question, a modality not supported by existing single-objective light-sheet techniques.

**Figure 1:**
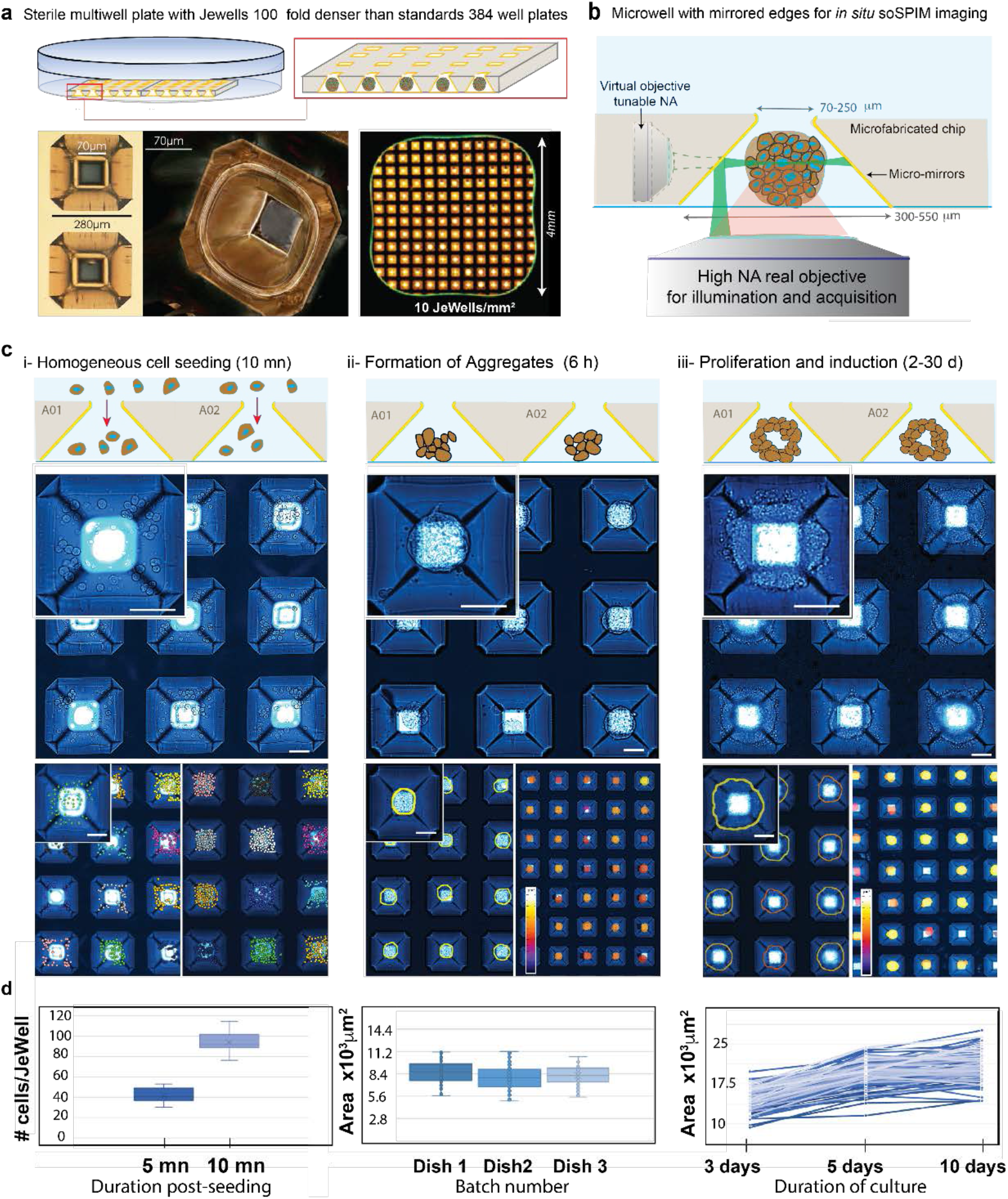
Principles of organoid culture in Jewells. **a**. Schematic of the organization of the Jewell array in a Petri dish. Close-up image of the JeWells inverted pyramid microcavities flanked with four 45° mirroring surfaces. The Jewells (golden) have a density 100-fold higher than a 384 well-plate (Green outline). **b**. Principles of the single objective light-sheet imaging (soSPIM). The 45° mirrors surrounding the organoids in the Jewells are used to reflect the excitation beam and create a light-sheet perpendicular to the optical axis. They act as a virtual objective. The emission of fluorescence following the light-sheet excitation is collected by the same objective used to create the light-sheet. **c**. Seeding and growth procedures in the JeWells chips. *i* - Dissociated cells are seeded in a single pipetting step on top of the whole Jewell array. The number of cells in each well is determined by the cell seeding density and fine-tuned by the incubation time before rinsing (5-10 mn). The initial number of cells is determined by bright-field microscopy. *ii-*Cells are left to aggregate for 6 hours and the initial size of each aggregate is measured. *iii*-Organoids grow inside the JeWells over prolonged periods. Growth, differentiation and imaging are all performed inside the same chip with a unique identifier for each organoid. The inverted pyramidal shape of the Jewell prevents material loss during all the pipetting and transport steps. All scale bars represent 50 µm. d. Quantification of the images above (N=100 JeWells for all panel).

### Production of calibrated organoids with no material loss

As a proof of concept, we studied the case of robust and fast developing neuroectoderm organoids generated from human embryonic stem cells (hESC). We used an eight days differentiation protocol to create neuro-ectoderm organoids^16^. We also tested our chips on hiPSC, primary hepatocytes (rat and human) with no noticeable differences as compared to hESC. We seeded dissociated hESC on top of the chip containing JeWells arrays. Buoyancy mismatch leads cells to sink at the bottom of the JeWells (**Figure 1-c**). After the seeding phase, the JeWells chips are rapidly screened in a phase contrast microscope, and the number of cells in individual Jewells are measured as a reference point (Online Methods). **Figure 1-d** illustrates the reproducibility and homogeneity of the robot-free seeding procedure. The initial cell concentration and the seeding time allows to precisely control the number of cells per well (eg: 0,5.10^6^ cells/ml for 10 min leads to 96 ± 10 SD cells/JeWell (N=100 wells)). We then monitored with phase contrast microcopy the projected area of the compact aggregates that form within the first few hours post-seeding (Online Methods). We observed a batch to batch reproducibility of 10% (3 batches) and 25 % dispersion of inter-well aggregate size (N=100 organoids) (**Figure 1-d)**. Phase contrast imaging was used over a prolonged period of time (up to 30 days) during the differentiation protocol to monitor the proliferation rate of the organoids. Thanks to the JeWells chip array geometry, each JeWell could easily be assigned an individual identification number, allowing unambiguous retrieval and imaging of each organoid at every time point, despite the need for periodically displacing the culture vessels from the microscope to the bio-hood and the incubator for medium exchange and cell culture. At the end of the culture, 90% of pyramidal shape Jewells were filled with a single organoid (**Supplementary Figure 4**), and 100% of the organoids remained in the Jewells despite all the manipulation steps.

### 3D live monitoring of organoid development

The previous characterization of the 3D culture only required 2D phase contrast imaging as reference. Next, we evaluated our ability to monitor hundreds of organoids in parallel by 3D light-sheet microscopy. We used a standard inverted microscope equipped with a soSPIM module^15^ and a motorized stage, and performed multi-position 3D time-lapse imaging over up to 8 days, with no noticeable drift (**Online Methods**). We first used calcein (40 µM in culture medium) as a cyto-compatible live extracellular fluorescent dye to negatively stain the developing organoids (**Supplementary Figure 5-a**). The 3D acquisition of each organoid, comprising of 70 optical Z-sections, took only 2s. 3D time-lapse fluorescence light-sheet imaging of organoids (**Movie 1**), demonstrated that calcein uniformly infused all the organoids, providing homogeneous access to nutrients and soluble factors inside the organoids grown inside the pyramids. The calcein staining revealed small lumens in the stem cell aggregates as well as cell division (**Movie 1**). We subsequently added Alexa-647, a non-permeable dye at (40 µM) to the calcein solution. We leveraged on the high-speed acquisition capacity of the soSPIM to quantify the diffusion rate of Alexa647 in the stem cell aggregates (**Supplementary Figure 5-b**), both in the intercellular clefts and in the lumens. It provided us with an indirect quantifiable proxy of the degree of compaction of the stem cell aggregates as well as the degree of tightness of the lumen edges. It demonstrates that small compounds have access to the internal part of the organoid fully embedded with 100% of Matrigel in less than 15 min, and the capacity of our platform to monitor their diffusion live. Altogether, this provides the quantitative evidence that the relative confinement of Jewells does not preclude accessibility to nutrients or extracellular compounds.

We next followed the development of the neuroectoderm organoid using stem cells expressing life act GFP and Histone 2B-mCherry^17^ (**Figure 2.a**) (**Movie 2**). We monitored 100 organoids in HCS mode, capturing one z-stack of 70 optical plans every 15 minutes for 8 days, representing a total of 500 z-stacks per organoid acquired in 125h) (**Online Methods**). Due to memory constraints, we only saved the total development course of 20 organoids randomly chosen among the one hundred monitored (**Figure 2.b-c**). At day 8, we did not notice any differences in the number of cells per organoids or shape between the imaged *vs* non-imaged organoids, confirming that the phototoxicity was negligible. After 5 days, rosette-like structures, i.e. cells aligning in 3D around a central streak ^18,19,20,21^ appeared in the organoids. By back tracking the rosettes from day 5, we could follow the time course of the first steps of development based on local cell arrangements (**Figure 2.d, Movie 3**). To confirm that the organoids differentiated in neuroectoderm, we collected them alive by peeling the micro-mirrored membrane from the glass slide (**Figure 2.e**), and subsequently performed qRT-PCR on eleven neuroectoderm specific markers (**Online Methods**). **Figure 2.f** unambiguously demonstrates the differentiation into neuroectoderm, characterized by a 10-fold reduction in E-cad and NANOG, and 500-fold increase in Pax6 for example.

**Figure 2:**
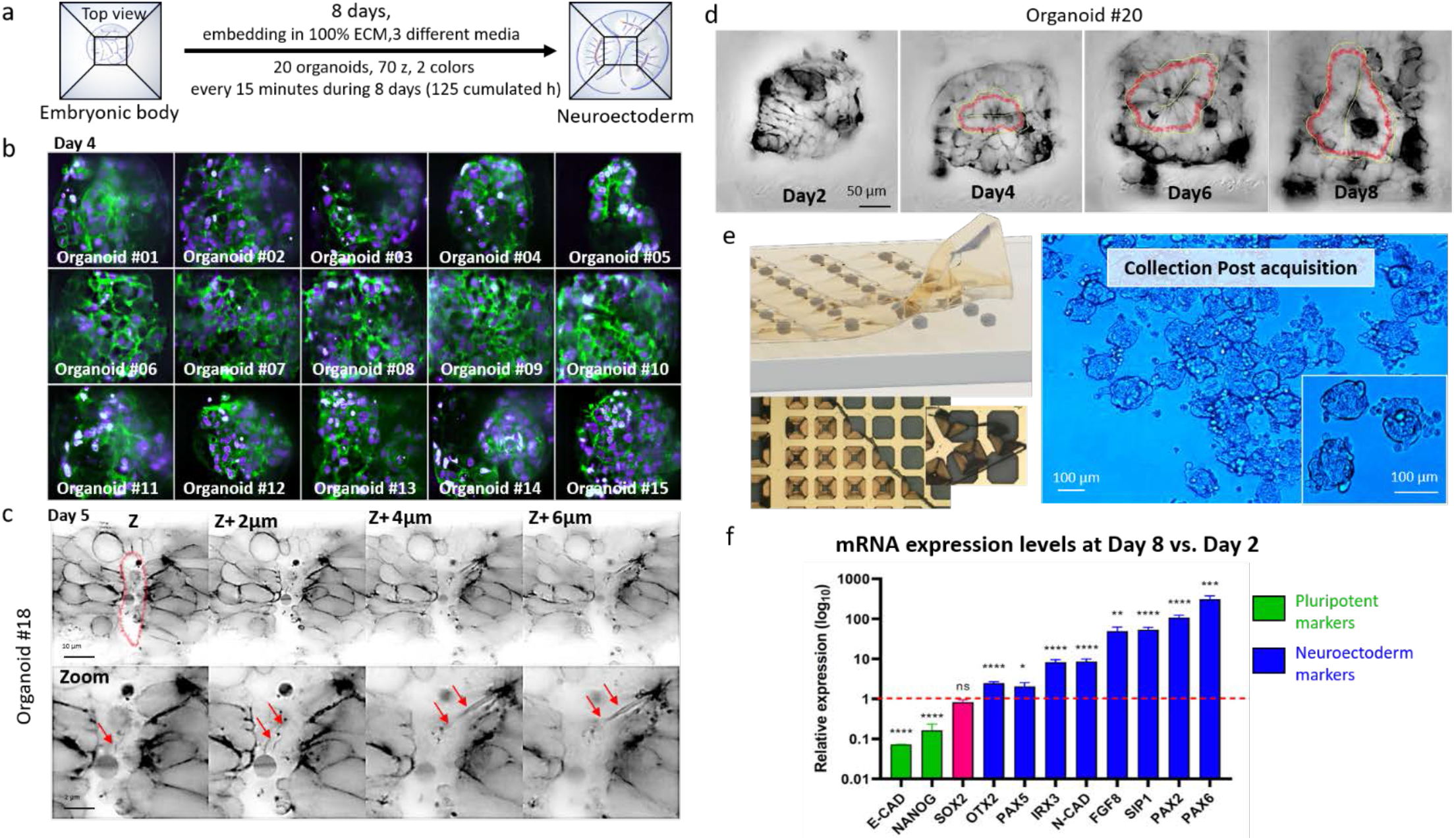
8 days 3D live imaging of stem cell differentiation into neuroectoderm. **a**. Summary of the experimental and imaging conditions: gallery of 20 organoids imaged sequentially in 3D (70 optical sections separated by 1 µm), every 15 minutes during 125 hours (spread over 8 consecutive days). **b**. Single optical sections gallery of 15 organoids expressing lifeact-GFP (green) and H2B-mcherry (histone, purple) 4 days after seeding. **c**. The high quality of the live soSPIM imaging of the lifeact-GFP signal allows to optically resolve the cellular arrangement and the cavitation events during the formation of a lumen (circled in red) in the middle of the organoids. Membrane protrusions (∼300nm) are also visible (red arrows). Lumenogenesis resulted in the expected creation of rosettes in the neuroectoderm organoids. **d**. Time lapse imaging example of the formation of rosettes (cells radially organized around a central streak) in 3D during the 8 days differentiation process. The rosettes are manually tracked (red contour) from the day of induction (day 2). **e**. Schematic of the Jewell chip peeling to collect the organoids after 3D live monitoring to enable mRNA extraction. **f**. RT-qPCR performed on 11 genes, demonstrating the expected reduction in pluripotent reporters and the increase of the neuroectoderm markers (N=3 replicates, > 1000 organoids/replicate) from the collected organoids. P-values: ****<0.0001, ***<0.001, **<0.01, *<0.05, ns: non-significative.

Taken together, these experiments demonstrated that our integrated platform is suitable for culturing organoids and 3D live imaging of them without phototoxicity, allowing continual assessment of their development. Moreover, retrieving the organoids after imaging enabled to determine their gene expression profiles or to collect them live for further use.

### High content 3D imaging of fixed organoids

We demonstrated the high content 3D imaging performance of our technique on fixed spheroids and organoids. As compared to commercial 96 well-plates (resp. 384), the large density of arrayed Jewells (10 JeWells/mm^2^) drastically reduces the total distance the translation stage needs to travel, from 89 cm (resp. 41cm) to 4.5mm to collect 96 organoids (**Figure 3.a**) and significantly reduces the evaporation and trailing of the immersion fluid. The specific geometry and organization of organoids in the JeWells chip allows their transmission image to be used as perfect beacons for the automated organoid selection and positioning of the light-sheet.

**Figure 3:**
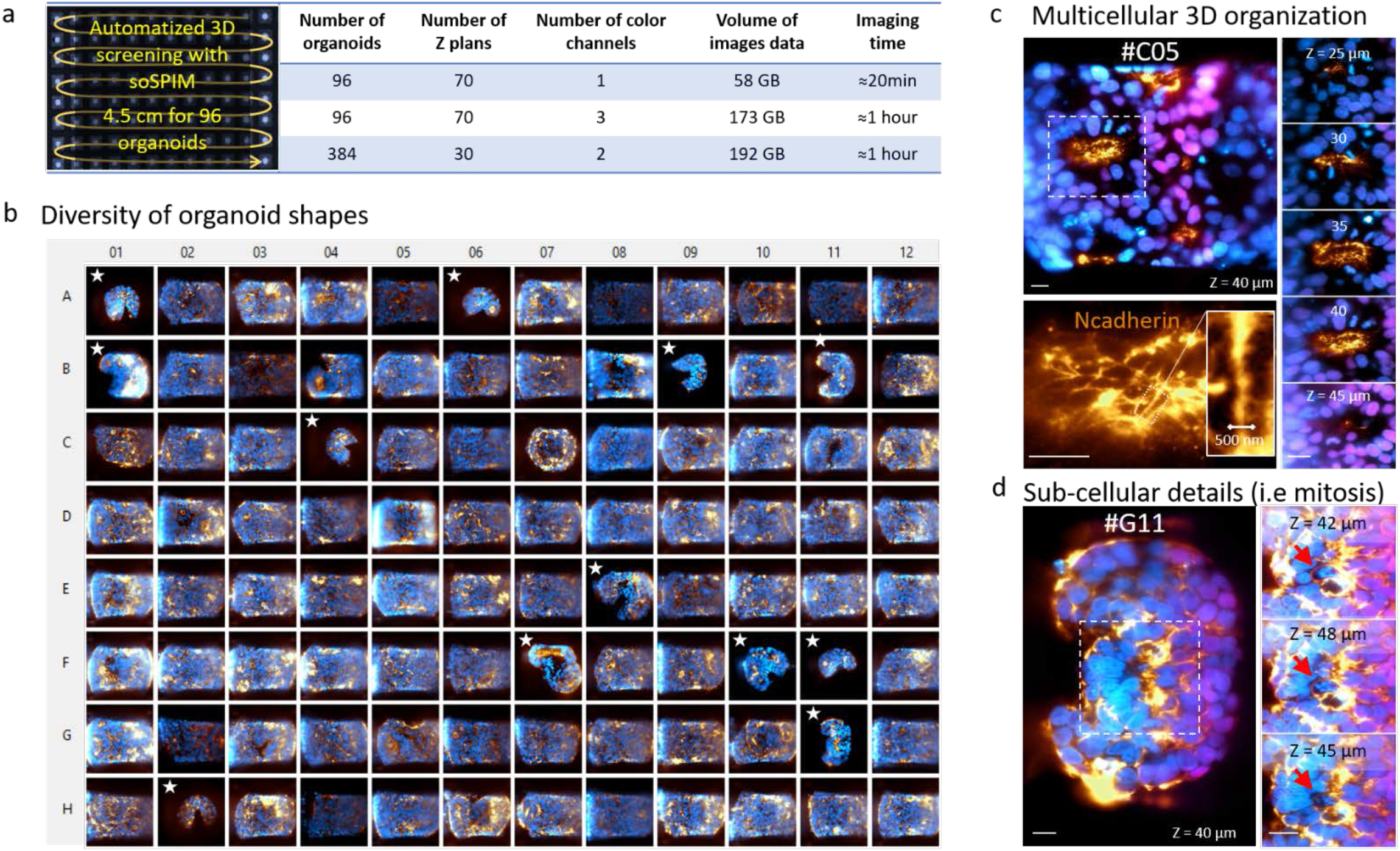
Specifications of the high content imaging platform for fixed organoids. **a**. Only 4.5 cm long path suffices to screen 96 well-arrayed Jewells containing organoids in a Jewell chip. The table summarizes the acquisition specifications in experimental imaging conditions (100 ms exposure time per plane, per color). **b**. 96 well-plate-like gallery of 96 3D organoids labelled with N-cadherin (gold), Sox2 (magenta) and DAPI (blue). The 96 organoids are a subset of 400 organoids acquired in an automatic single workflow using our HCS platform. Organoids with B-shapes (marked by white stars) and the O-shapes have been identified as two morpho-classes for the organoid shapes in the culture. **c**. Representative 3D high-resolution images of internal multicellular organization of a rosette core of an O-shape organoid. The N-cadherin staining highlighting the complex geometry of the filaments inside the streaks. **d**. Representative 3D high-resolution images of sub-cellular structures (i.e condensed chromosomes of a mitosis, red arrows) observed in the central region of a B-shape organoid.

We fixed the neuroectoderm organoids at day 8 (Online Methods). The pyramidal shape of the Jewells prevented any material loss during the 30 steps of pipetting and solution exchange required to complete the staining procedure. Setting-up the entire screening parameters and selection of 96 organoids took around 10 min (at 60x magnification), including the high-resolution pre-screening of the JeWells chip. The time between the acquisition of two adjacent Jewells was 1.5s and the stability of the light-sheet thickness below 3%. We performed one side light-sheet illumination and used a 60x WI 1.27NA objective for the illumination and the collection of fluorescence. Each 3D imaging of immunostained fixed organoids took about 7s per wavelength for 70 optical sections.

**Figure 3c** displays a gallery of 96 organoids acquired with 3 wavelengths (Nuclei (405nm), Sox2 (488nm) and N-cadherin (647nm)). Each z-stack composed of 70 optical sections in 3 colors was acquired in 21s. It took 1 hour to acquire the 96 organoids in 3D with 3 colors, representing 173GB of raw data (**Figure 3.a** for benchmarking). We reached a spatial resolution of around 300 nm in XY. The light sheet thickness was 3.7 μm wide at the thinnest part for a length of 60 µm (**Supplementary Figure 1.c**). We created libraries of 100 to 400 organoids images in 3D of which we grouped in 96 well-plate like format for display purpose (**Figure 3.b**). The spatial resolution was sufficient to clearly detect rosettes in 3D (**Figure 3.c**), nucleoli and mitosis (**Figure 3.d**), and actin rich protrusions (**Figure 3.c-d**) in 200 μm diameter organoids. Single-sided light-sheet illumination provided an expected sectioning gradient along the propagation direction of the light-sheet (due to scattering and Gaussian beam illumination). However, we also implemented a multi-angle light-sheet illumination mode that leverages the arrangement of the 4 side-mirrors of the JeWells containing organoids (**Movie 4-5**). In the context of HCS, we found no real benefit of multi-side illumination, since it is more time-consuming, photon-demanding, and requires dedicated image fusion algorithms. Another benefit of the stereotypical positioning of the organoids inside the wells was that it made any post processing detection of organoid position unnecessary.

**Figure 3.b** illustrates the variability of organoid shapes in a subset of 96 organoids, which we divided in two morpho-classes: bean shaped (B-shaped) and spherical (O-shaped) organoids. The native quality of the 3D images allowed multicellular structures, called neuroectoderm rosettes, to be clearly identified (**Figure 3.c**). Moreover, the spatial resolution and contrast of the images enabled unambiguous visualization of chromosome condensation during mitosis in the core of the organoids (**Figure 3.d**).

### Multiscale analysis of morphogenetic parameters

Beyond visual inspection of the organoids, the high-resolution of the soSPIM images allows further automatic quantification. We developed automated segmentation and classification techniques using traditional and deep learning-based image analysis at whole organoid, multicellular, and subcellular scales. First, we used a custom-made segmentation routine based on local thresholding and 3D water-shedding (**Online Methods**) to segment nuclei (**Figure 4-a**). We computed the mean intensity for each nucleus to quantify the individual expression levels of transcription factors (such as Sox2). This allows automatic evaluation of different cell populations within the organoid gated by their expression level.

**Figure 4:**
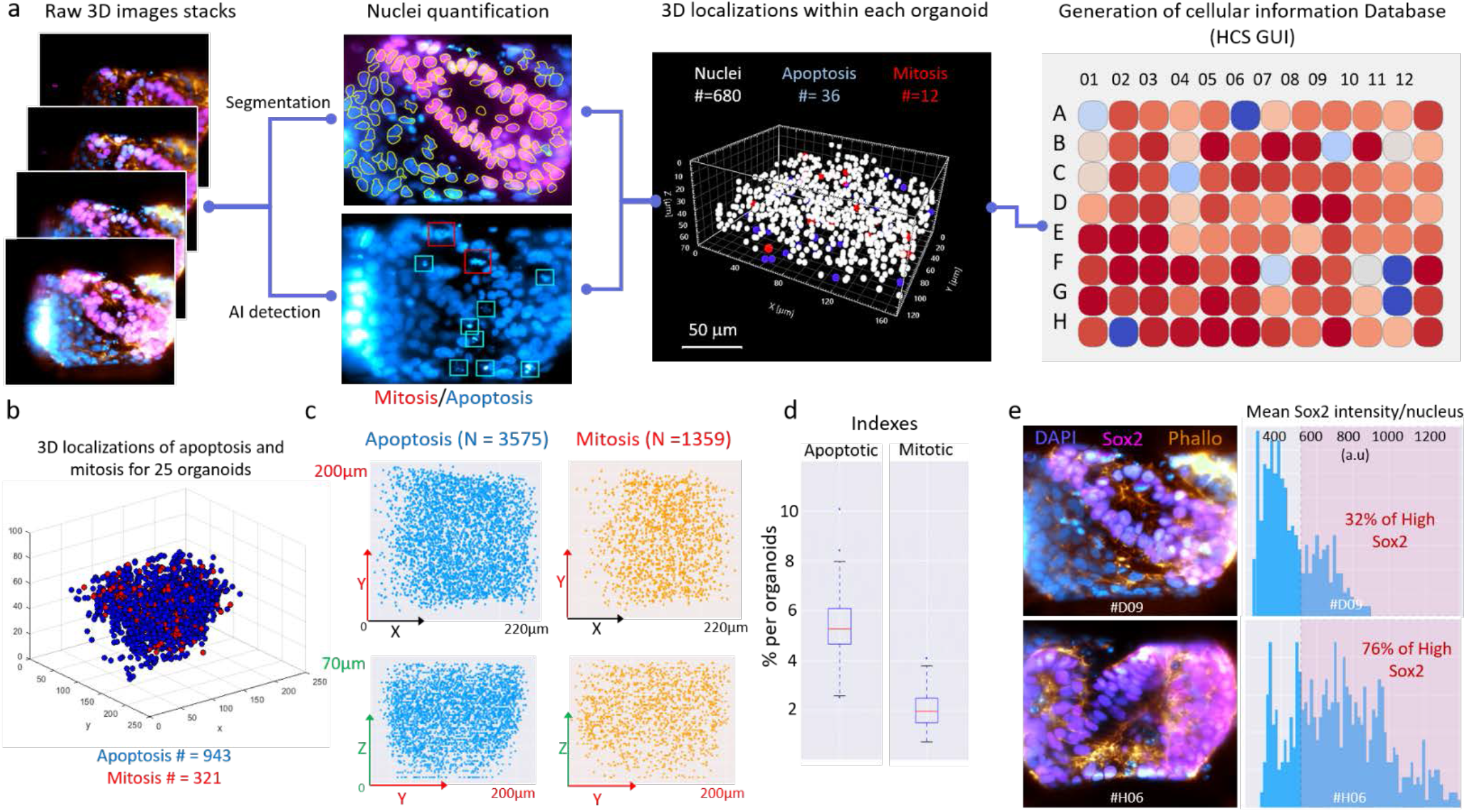
Cellular events quantification in organoids. **a**. Image analysis workflow for automated nuclei segmentation and deep learning-based detection of mitosis and apoptosis in individual organoids. Central image shows an example of a 3D localization map of all nuclei (N=680), apoptotic bodies (N=36) and mitosis (N=12) of one organoid. All these quantitative features are organized in a database for datamining using CellProfiler Analyst software. Each box in the heatmap (A01, A02,…, B12) corresponds to a single organoid. The color-code accounts for the number of nuclei for each organoid. **b**. The self-centering localization of organoids in JeWells enables to generate the cumulative 3D spatial distribution of all the apoptosis (blue dots) and mitosis (red orange dots). N=25 organoids shown. **c**. 2D projection of the apoptosis (blue) and mitosis (orange) distributions for the 96 organoids, in the lateral (XY) and axial (XZ) planes. **d**. Proportion (%) of mitosis and apoptosis relative to the number of nuclei. N = 96 organoids. **e**. Representative images of organoids with low (top) and high (bottom) numbers of Sox2 ^+^ cells. Corresponding distribution of nuclear expression levels of Sox2 for each segmented nucleus in the organoid and nuclei percentages with ‘High Sox2’ level.

Then, we leveraged the large number of high-resolution 3D images to perform deep learning segmentation of apoptosis and mitosis events. We trained a convolutional neural network YOLOv2 from a subset of 28 organoids, by manually creating standardized bounding boxes around the events (1123 apoptosis and 344 mitosis events) (**Online Methods, Supplementary Figure 6.a**). It enabled the automatic localization of apoptosis and mitosis events in the organoid 3D images, with 89% accuracy (4% false positive, 7% false negative, benchmark performed on 20 organoids by a trained human). We performed 3D mapping of the positions of the nuclei, mitotic and apoptotic events to each organoid (**Figure 4-a**), which we organized in a database for display by CellProfiler Analyst, a cytometry-like freeware^22^ (**Figure 4-a**).

The cumulative spatial distribution of the mitosis and apoptosis events for the 96 organoids (**Figure 4b-c**) displayed no preferred loci for apoptosis or mitosis in the organoid volume. This confirms that Jewells do not bias the organoids development. We quantified that 5± 2.9 % of the cells are apoptotic and 2±1 % are mitotic (**Figure 4-d**). The expression levels of Sox2 was spatially heterogeneous in the organoids. The bimodal distribution of expression levels at the single nuclei level quantifies the proportion of Sox2- and Sox2+ cells in each of the 96 organoids (**Figure 4-e**).

We then trained (**Online Methods**) the shape recognition Convolutional Neural Network (CNN) DenseNet121^23^ to identify automatically the 2 morpho-classes of B and O shape organoids (**Figure 5-a**). Over 96 organoids, we detected 12 B-shapes, which is 99 % in agreement with human verification (**Supplementary Figure 6-b**). In comparison with O-shape organoids, B-shape organoids systematically displayed fewer nuclei and no internal rosette or streaks but showed similar distributions of Sox2 positive cells.

**Figure 5:**
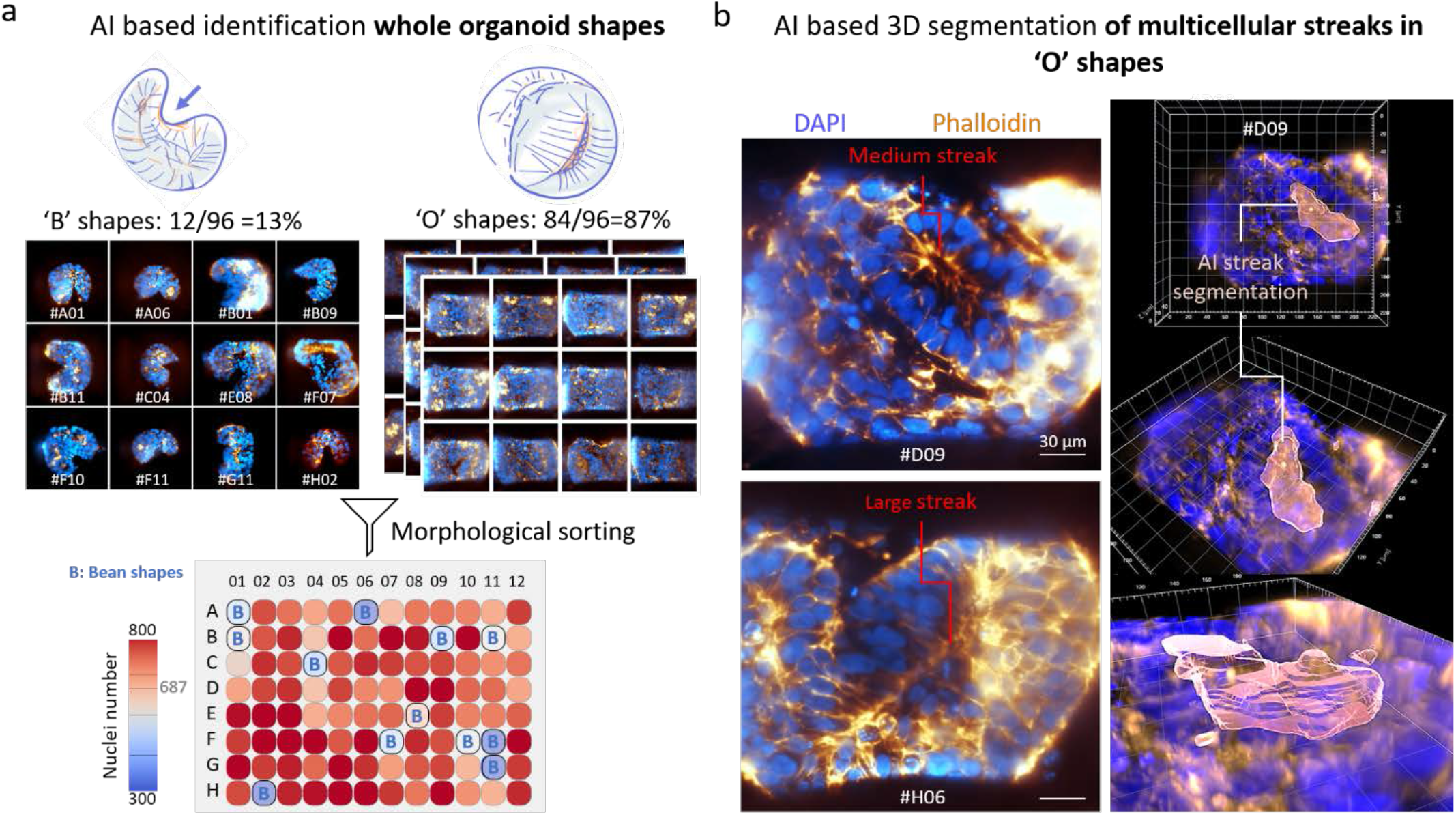
Quantification of morpho-descriptors for organoid shapes and multicellular streaks. **a**. Automatic classification of the neuroectoderm organoids morpho-classes (B-shape or O-shape) using Convolutional Neural Network (CNN) DenseNet121 for 96 organoids. The attribution of a ‘B’ or ‘O’ tag to each organoid in the database was successful with 100% accuracy. We excluded the non-rosette presenting B-shaped organoids for the subsequent analysis. Color-code of the heatmap represents the number of nuclei per organoid, grey value 687 is the mean value for the 96 organoids. **b**. Two examples of rosette 3D streaks inside O-shaped organoids. Streak regions based on actin structure (Gold) were automatically segmented using 3D U-net Neural Network. Subsequent thresholding of the prediction probability map enabled the automated 3D segmentation of the streaks.

Lastly, the automated segmentation of the individual rosette streaks inside the O-shaped organoids was obtained by training a 3D U-net deep-learning algorithm from the 3D actin images (**Online Methods, Figure 5-b, Movie 6-7**). We extracted in the database 415 relevant streaks longer than 2 cells (20µm), from which we computed the morphological descriptors detailed in **Supplementary Table 2**.

The final database hence contained multidimensional information, at the cellular (nuclei, apoptosis, mitosis), multicellular (rosettes morphologies) and whole organoid (shape) levels (**Figure 6.a**). It provided the quantitative description of the organoid culture that could serve to assess the quality of the organoid development or correlate with the expression of growth factors and morphological descriptors. The literature reports prominent recruitment of Sox2+ cells around rosette streaks in 2D^18,19^ and in 3D^20,21^. Our database provides a quantitative view of the localization of Sox2 expression along the streaks. The scatter plot of the rosette streak volumes vs the average Sox2 expression levels per nucleus of each organoid (**Figure 6.b**) can be gated for large (+) or small (-) volume and expression levels. It defined 4 populations referred to by +/+, +/-, -/+ and -/- (streak volume/Sox2 expression). Organoid clustering in the +/+ quadrant of the plot show that large streaks (volume >3.10^4^μm^3^) unambiguously displayed a high number of Sox2 positive cells around them. Relating the multiscale descriptors contained in the database (**Figure 6 b**), we showed a significant increase in the total numbers of cells and of mitosis in the +/+ *vs* -/-population. The scatter plot also shows that small streaks, can exist in organoids either with very few Sox2+ cells (-/-quadrant), or very large number of Sox2+ cells (-/+). It clearly indicates that the expression of Sox2 is not causal, nor a *bona fide* reporter of streak organization.

**Figure 6:**
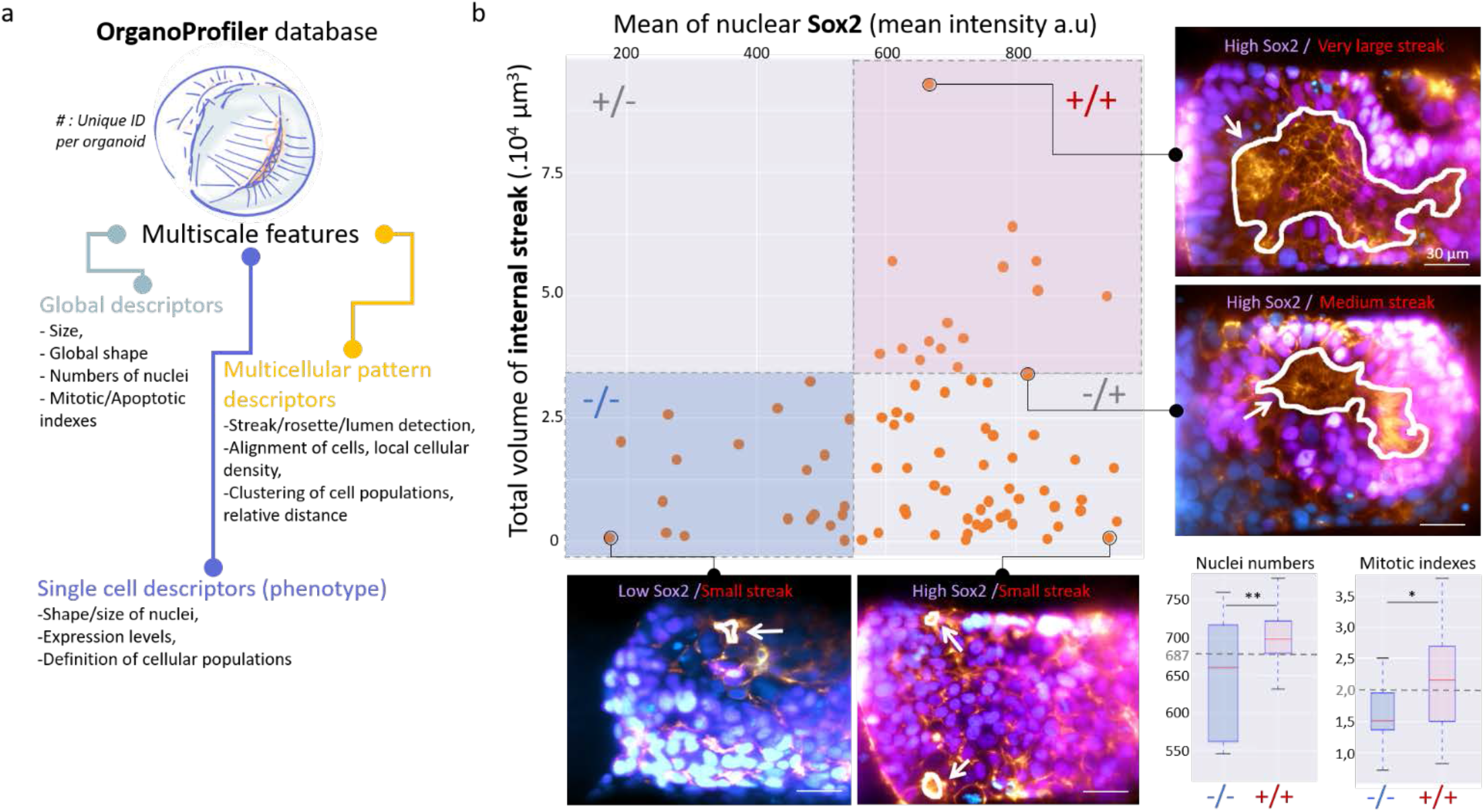
Datamining of the organoid database. **a**. Schematic representation of the multiscale information quantified from the 3D images of the organoids. **b**. Scatter plot of the rosette streak volumes *vs* the average Sox2 expression levels per nucleus, for each O-shape organoid. Each individual point represents one organoid. The populations were classified in 4 groups using gating based on the Sox2 intensity as described in Figure 4-e and Online Methods. Images show examples of organoids in each of the 4 groups. White lines represent the AI-based segmentation of the streak contours. Box plots represent the distributions of the number of nuclei and mitotic indexes for the -/- and +/+ gated populations. ** : p-value < 0.01; * : p-value < 0.1. Grey values (687 and 2,0) represents the mean values for the entire O-shape populations of organoids.

## DISCUSSION

Our integrated high content imaging platform combines the advantages of high-density organoid culture with light-sheet-based 3D fluorescence microscopy. It allows monitoring in 3D hundreds of live organoids with an acquisition rate of 300 organoids per hour, in a fully automatic manner and with no apparent phototoxicity. In the present work, we used organoids of 150 to 200 μm in diameter. Light-sheet illumination allows a typical penetration depth ranging between 60 to 100 μm in live samples, which is an inherent limitation of single photon imaging across diffusive samples. We used corresponding JeWells cavities with 70 μm aperture and 290 μm base, but Jewell dimensions can be scaled up. The maximum lateral size is restricted by the field of view of the objectives (25mm for Nikon at the image plane), since soSPIM requires both the mirror and the samples to fit the field of view. Therefore, with a 60x magnification we are limited to 200 μm width organoids, and with a 20x magnification to 600 μm width organoids. We also note that the Jewells used here have a square shape but can be easily turned into rectangular grove to grow elongated organoids. This dimension limit is not really an issue since most reported organoids are within this size range: cancer pancreatic^24^: 100 μm, intestinal^25^: 50-200 μm, gastruloids^26^: 100 μm diameter and 400 μm length, liver^27^: 300 μm, micro kidney^28^: 150-200 μm, cancer spheroids^29^: 250-750 μm.

JeWells culture vessels can also be used to grow different types of cells. For example, we successfully cultured primary and hiPSC derived hepatocytes (**Movie 8**). We also performed a 3D time-lapse experiment of living HE3B cancer cell spheroids stably expressing H2B-GFP. We could image them every 90 seconds during 18h, representing 1.23 TB of raw data, without visible signs of phototoxicity or photobleaching (**Supplementary Figure 3**). Such acquisition rate was not possible on a spinning disc microscope.

A direct comparison of our high content imaging platform with a spinning-disk-based commercial HCS platform showed improved acquisition speed, increased penetration depth, and a strong reduction in laser radiation (**Supplementary Figure 3**). As an illustration, we could acquire 96 3D organoids in 3 colors in only 1h, versus 8h for the Opera system.

In contrast with other single objective SPIM approaches dedicated to organoid investigations, our approach streamlines 3D cell culture and high content live imaging with no sample manipulation, quantitative image analysis, and datamining. It can be easily mounted on any standard inverted microscope, after simple upgrade. The large number of organoids that can be monitored simultaneously allows for training of convolutional neural network for deep learning analysis, opening great promises for the predictive analysis of growing organoids by artificial intelligence. As a simple illustration, we demonstrated our approach to allow extracting meaningful quantitative information at various spatial scales using existing deep learning algorithms, directly from the raw data.

Assessing the quality of organoid cultures as they grow is a real challenge. It is currently performed visually at their final stage by the expert eyes of trained biologists, making the overall process fastidious. We here established that the technique can quantify generic multiscale features such as mitosis, apoptosis, cell organization and whole organoid shape as well as follow the live development of 3D cell culture. We think our approach, combined with dedicated artificial intelligence, has the promising potential to provide non-invasive quality control and guidance in the establishment of differentiation protocols. Before this becomes a reality, we envisage sharing our libraries of 3D organoid images to prompt the development of AI-based quantitative analysis and quantitative benchmark of organoids culture. The libraries of the 500 organoids used in this study will be available on the Mechanobiology Institute Server. Altogether, we think this work can boost organoid research to reach its full potential.

## DATA AVAILABILITY

Raw data of the 500 organoids acquired in this study will be available on the server upon request We will provide free access to JeWells chips for testing based on Material Transfer Agreement.

## ACKNOWLEDGMENTS

VV acknowledges the support of NRF investigator, MBI seed funding and ANR. JBS and RG are supported by the Labex BRAIN, Idex Bordeaux, ANR soLIVE and France BioImaging infrastructure (ANR-10-INSB-04). CNRS PhD fellowship for TD. We thank F. Saltel (BaRiTon lab) for providing the HEP3B-H2B-GFP stable cell-line. We acknowledge the kind gift of Lifeact/H2D Esc from O.Reiner (Weizmann) and C. Butler for discussions. A.B and GG acknowledges the support from MBI core funding. All authors thank A. Wong and D. Pitta de Araujo for his help editing the manuscript.

## ONLINE METHODS

## Online Methods

### Fabrication methods for the Jewells chips

Jewells pyramidal microwells are produced by mean of a casting procedure which uses capillarity filling to form a textured polymeric thin film. The textured film is then coated with metal to enhance the reflectivity of the micro-mirrors and finally glued to a coverslip. A polydimethylsiloxane (PDMS) mold with an array of pyramidal micro-pillars is prepared as a cast-replica out of a silicon primary mold (**Supplementary Figure 1a**). The silicon primary mold is made in according with a protocol published by Galland et al. ^1^. PDMS Sylgard 184 (Dow Corning) base resin and reticulation agents are mixed in 10:1 ratio and out-gassed (20-30 min in a vacuum jar evacuated to 5-10 mbar). The liquid PDMS is poured on the silicon mold and further out-gassing ensure filling of the cavities. (vacuum jar at 5-10 mbar for 10 min). After thermal curing (2h at 75 °C on a hot plate) the PDMS mold is peeled-off and cut in 1 x 1 cm^2^ pieces. Each 1 x 1 cm^2^ cut can be used multiple times to produce Jewells devices.

A PDMS 1×1 cm^2^ cut is flipped face-down on a flat PDMS substrate (**Supplementary Figure 1a**), then a small amount of liquid UV-curable optical resin (NOA73, Norland Optics, USA) is dropped at one edge of the mold. Driven by capillarity, the liquid fills the cavity between the PDMS mold and the flat substrate. Once the whole structured cavity is filled, the NOA is cured with flood exposure to UV light (UV LED KUB2 light, Kloe France, 365 nm with power density of 25 mW/cm^2^, 1 min exposure). After UV-curing, the 1 x 1 cm^2^ PDMS mold is removed leaving the cured NOA on the flat substrate. Excess material is trimmed on all 4 edges and a film of Au is deposited by sputter-coating (Jeol Auto fine-coater, Jeol Japan. A process with setting 20 mA, 45 s is repeated 4 times to achieve a final thickness of the coating of about 40 nm, with 1 min waiting between each step to allow for cooling. Meanwhile, a glass coverslip (such as #1.5H) is prepared as a final substrate for the Jewells device by coating it with a thin layer of NOA73; using spin coating as the method of deposition, about 20-30 µm thick NOA is deposited on the coverslip and pre-cured under UV-light (30s in the same UV LED KUB2) so to become solid but still retaining adhesive properties. The Au-coated NOA structured film (still on the flat PDMS substrate) is flipped and placed in contact with the NOA-coated coverslip and gently pressed to ensure good contact. The NOA adhesive layer is fully cured with a final step of UV flood exposure (1 min in the same UV LED KUB2 as previously) the flat PDMS substrate can be peeled-off leaving removing at the same time portion of the metal film which was in direct contact with the PDMS. Finally, the Jewells device is ready, with top base of the pyramidal cavities open thanks for the Au removal with the flat PDMS substrate.

### Long term passivation of JeWells chips

This step is mandatory to avoid any adhesion of cells to the chips during the time of the experiments (>25 days) and to allow 3D culture of the cells. A solution of Lipidure® (CM5206, NOF America) at 0.5% (w/v) in pure ethanol is added to completely cover the Jewells chips. Optionally, a plasma cleaning of the coverslip (5min) has be done before. The Jewells chips covered with Lipidure solution is out-gassed (5-10 min in a vacuum jar evacuated to 5-10 mbar. Then, the ethanol-Lipidure solution is allowed to completely be evaporated under a sterile hood (>2 hours) in order to form a coating with same molecular structure as polar group of phospholipid. Before cells seeding, Jewells chips are rinsed with PBS or cell culture medium and out-gassed as previously.

### Stem cell lines

The human embryonic stem cell lines used for the experiments were H1 and K21 (lifeact-GFP, H2B-mCherry) that were obtained from WiCell Research institute, Inc. (Madison, WI, USA) and a gift from Orly Reiner’s lab (Weizmann Institute, Israel), respectively. Both cell lines used were on early passage number and confirmed to be contamination free.

### Maintenance of human pluripotent stem cells

The basement membrane matrix used to culture and maintain the cells was hPSC-qualified Matrigel^™^ (354277, BD Biosciences) and the maintenance medium was mTeSR^™^1 medium (05850, StemCell^™^ Technologies) for H1 and RSeT™ medium for K21 (05978, StemCell^™^ Technologies). To passage the cells, the 80% confluent hPSC culture was enzymatically treated with Dispase (07923, StemCell^™^ Technologies) for H1 and TrypLE™ Express Enzyme (12605028, Thermofisher scientific) for K21 followed by mechanical scrapping and suspending in fresh medium to re-plate on matrigel coated wells.

### Embryoid body formation in Jewells

For formation of embryoid bodies (EB) in Jewells, a single cell suspension of hPSCs was generated by enzymatic treatment of H1 and K21 cells with ReLeSR™ (05872, StemCell^™^ Technologies) for about 9 minutes. The single hPSC cells were suspended at a concentration of 0.5 million cells/ml in EB formation medium supplied in STEMdiff™ Cerebral Organoid Kit (08570, StemCell^™^ Technologies) supplemented with 10 μM Y27632 (72304, Calbiochem, Merck Millipore) and seeded on to Jewells. The cells were incubated for 15 minutes to allow adequate number of cells to enter per Jewells. Then the Jewells were washed with DMEM/F12 medium to remove excess cells and incubated in EB formation medium supplemented with 10μM Y27632 for 24 hours. The neuroectoderm differentiation of EBs in Jewells was then initiated after 24 hours of cell seeding and/or when the EBs where ≥ 120µm in size.

### Induction of neuroectoderm in hPSCs

To induce neuroectoderm differentiation, the embryoid bodies were cultured in STEMdiff™ Cerebral Organoid Kit (08570, StemCell^™^ Technologies) as per manufacturer’s instructions with slight modifications. Briefly, when the EBs in Jewells reached ≥ 120µm in size they were induced to neuroectoderm using induction medium from Cerebral Organoid Kit for 4 days. On day 5, induction medium was removed and the induced EBs were embedded in 100% hPSC-qualified Matrigel^™^ (354277, BD Biosciences) for 30 mins. Once the induced EB in Jewells were coated with a layer of matrigel, they were allowed to undergo neural expansion using expansion medium from Cerebral Organoid Kit for 3 days.

### Primary rat hepatocytes isolation and seeding in JeWells

Hepatocytes were harvested from male Wistar rats by the two-step collagenase perfusion method as described by Seglen et al.^2^. Animals were handled according to the protocol established by Institutional Animal Care and Use Committee (IACUC) approved by the IACUC committee of National University of Singapore. Each isolation yielded >108 cells, with the viability ranging from 75-90% as tested by Trypan Blue exclusion assay. 0.15×106 cells/ml suspended in cold hepatocytes culture medium (William’s E media (A1217601,Thermofisher scientific) supplemented with 1 mg/ml BSA (A7906,Sigma-Aldrich), 100nM Dexamethasone (D4902,Sigma-Aldrich), 2mM L-Glutamine (25030081,Thermofisher scientific), 0.3μg/ml of insulin (I9278,Sigma-Aldrich), 50μg/ml linoleic acid (L1376,Sigma-Aldrich), 100 units/ml penicillin and 100μg/ml streptomycin (15140122, Thermofisher scientific) were seeded as previously described per JeWells dish and the dishes placed in the incubator for 5-10min to allow for the cells to settle. The dishes were then swirled, and the supernatant was collected to remove the floating cells. 1ml of fresh hepatocytes culture medium was added to the dish and were left to incubate during 2 to 4 days before fixation and immunostaining.

### Cancer cell lines seeding in JeWells and formation of spheroids

HEP3B, an hepatocellular carcinoma cell line, modified to express H2B-GFP (gift from F. Saltel BaRITOn Lab, Univ. Bordeaux, INSERM UMR1053) were suspended at a concentration of 0.5 million cells/ml in growth medium (DMEM (biowest, L0106) supplemented with 10% FBS (Sigma), 1% GlutaMax (Sigma), 1% Penicillin-Streptomycin (Sigma)) and seeded (1ml) on top of a JeWells Chips. After 10 min, supernatant has been replaced to remove floating cells and spheroids were allowed to grow during 36 to 48h before fixation and immunostaining.

### JeWells Peeling and Real-Time PCR

The organoids were released from the Jewells by peeling off the top layer. The wells were then flushed twice with DMEM/F-12 1:1 (Gibco) and the released organoids were collected on a microcentrifuge tube. Total RNA was extracted from these organoids using RNeasy Plus Micro Kit (74034, QIAGEN) as per manufacturer’s protocol. 450ng of total RNA was used to form the cDNA using cDNA synthesis kit (SensiFAST™, BIO-65054, Bioline). qPCR was performed using FastStart Universal SYBR Green Master (ROX) mix on a CFX96 Touch Real-Time PCR detection system (Bio-Rad). Primers sequences for the 12 genes tested are listed on the Table 1. GAPDH was used as a housekeeping gene.

### Immunofluorescence staining for imaging

Organoids/spheroids in the Jewells chip were fixed for 20 minutes in 4% paraformaldehyde (28906, Thermofisher scientific) at room temperature, and permeabilized for 24 hours in 1% Triton X-100 (T9284, Sigma Aldrich) solution in sterile PBS at 4°C on an orbital shaker followed by 24 hours incubation in blocking buffer (2% bovine serum albumin BSA (37525, Thermofisher scientific) and 1% Triton X-100 in sterile PBS) at 4°C on an orbital shaker. After overnight incubation, samples were incubated with primary antibodies anti-SOX2 (goat, AF2018, R&D Systems, 1:50), anti-N-cadherin (mouse, ab19348, Abcam, 1:100), or anti-MRP2 (mouse, M2III-6, NBP1-42349, Novus Biologicals,1:100) diluted in antibody dilution buffer (2% BSA, 0.2% Triton-X 100 in sterile PBS) at 4°C for 48 hours. The samples were then rinsed 3 times with washing buffer on an orbital shaker (3% NaCl and 0.2% Triton-X 100 in sterile PBS) and then incubated with secondary antibodies, donkey anti-mouse IgG-Alexa Fluor® 488 (A21202, Invitrogen, 1:500), donkey anti-goat IgG-Alexa Fluor® 633 (A21245, Invitrogen, 1:500), 0.5μg/ml DAPI (62248, Thermofisher scientific) and Alexa Fluor® 488 phalloidin (A12379, Thermofisher scientific, 1:200) at 4°C for 24 hours on an orbital shaker followed by 5 steps of rinse with washing buffer. The samples were then washed following similar washing steps as after primary antibody incubation. After that, samples were mounted using RapiClear® 1.52 (RC 152001, Sunjin lab) pre-warmed to 37°c.

### Automated acquisition process

MetaMorph software (Molecular Device) was used to control the microscope, the (X,Y,Z) motorized stages, the soSPIM device and steer the high content screening through the *Multidimensional Data Acquisition (MDA)* module. A dedicated home-made plugin enabled to control the soSPIM beam steering unit, and to synchronize it during the 3D acquisition process, as described in Galland et al.^1^. The soSPIM specific optical architecture couples the lateral (XY) stage position with the axial (Z) position of the light-sheet after reflection onto the 45° mirror of the JeWells (**Figure S2-A**). While this coupling makes the 3D sectioning very easy by simply moving the light-sheet along the mirrors, it has the downside to induce unwanted axial drift of the light-sheet upon lateral drifts of the sample. To overcome this potential drift issue, we implemented a fully automatic positioning of the light-sheet based on JeWells bright-field images (**Figure S2.B**). It allows both to automatically position the light-sheet during the multi-well screening, and the automatic correction of long-term spatial drifts. The automatic positioning relies on the mathematical cross-correlation between the bright-field images of the JeWells acquired during the screening, with a reference image (**Figure S2.B**). This cross-correlation algorithm was integrated into the acquisition pipeline, and executed at different key time points during the screening. It returns the drift vector in calibrated units, and corrects for it by translating the XY stage with the opposite vector. We precisely evaluated the repositioning accuracy of our approach by randomly displacing the stage around its initial reference position, followed by the automatic repositioning (100 measurements, random positions between -3 µm and +3 µm in X and Y directions). As a readout of the capability of our system to maintain the light-sheet position in focus, we measured the light-sheet width in the focal plane of the objective, and computed its deviation from the initial reference value, before and after drift correction (**Figure S2.B**). We measured an average broadening of the light-sheet of 0.085 +/-0.033 µm, which represents a fluctuation of less than 3% for a 2.5 µm thick light-sheet, as used in our high content acquisition ensuring optimal 3D imaging.

Optimal high content imaging was performed using the dense well-arrayed JeWells containing organoids. It is a 4-steps process as described in **Figure S2.C**. Briefly, in *<u>step 1</u>*, a large field preview of the chip (of 8 x 8.8 mm for 300 JeWells) is automatically acquired in bright-field and epi-fluorescence modes using the MetaMorph *ScanSlide* module (**Figure S2.D**). *<u>Step 2</u>*: all the JeWells are automatically detected from the preview bright-field image using a home-made macro based on Fourier-filtering and automatic thresholding. *<u>Step 3</u>*: once detected, it is possible to quickly and interactively manually select/unselect the positions to acquire, which will be directly loaded into the *MDA* module (**Figure S2.D**). *<u>Step 4</u>*: the high content imaging sequence is launched, and the recorded positions are automatically screened following the acquisition parameters preliminary defined (eg: time-lapse frequency, 3D stack definition, number and selection of excitation wavelengths). During the screening acquisition process, the JeWells positions are automatically updated by cross-correlation, at the first time-point, and every *N* time-points (*N* is a parameter that can be adjusted depending on each system stability, which we adjust to correspond to a repositioning every 30 min in our microscope) ensuring optimal 3D imaging over the whole acquisition process duration. Overall, it took 25 min to setup the HCS experiment containing 321 positions (*steps 1* to *3*) including 20 min for the chip preview acquisition. For 96 organoid acquisition, the setup time was reduced to 10 min.

### Nuclei segmentation

Nuclei acquired with the soSPIM were segmented using a custom-made macro developed in imageJ/Fiji. Briefly, raw images of nuclei (DAPI) were binarized plan by plan by local thresholding with same parameters for all organoids (plugin of G .Landini, Niblack method^3^, radius of 50 pixels) followed by closing and holes filling morphologic filters. Finally, 3D watershed was performed in 3D using the MorphoLibJ plugin^4^ to separate touching nuclei. Segmentation applied to 96 organoids resulted in 65999 nuclei and has been used to measure intensities of transcription factors for all organoids. The nuclei located at the edges of the light sheet were excluded from the detection to minimize any biased. To review of the detection quality, we manually assessed the error of the whole process. We found that 85% of the nuclei were properly detected. This intrinsic error is accounted for in our error bars.

### Deep learning-based detection of mitosis and apoptosis in 3D

The detection of apoptosis and mitosis events was performed using YOLOv2, the convolutional neural network published by ZeroCostDL4Mic. We used the default settings. We run it on our own GPU resources (NVIDIA Quadro RTX6000 24GB) (**Supplementary Figure 6a**). We manually selected 30×30 pixels bounding boxes around the central plane of the apoptosis and 50×50 pixels bounding boxes around the central plane of the mitosis, on 28 organoids with 71 z-planes using ImageJ. We used the DAPI channel. These annotated stacks constituted the ground truth of the training dataset. We selected a total of 344 mitosis and 1,123 apoptosis. We extracted from each event z-plans corresponding to the centre slice bounding box as well as one slice above and one below. Using image flipping and rotation we finally augmented the training set by 4 folds. In the end the neural network as trained on a final dataset of 2D vignettes representing 4,128 mitosis and 13,476 apoptosis. We used 90% of the data set for training and 10% for validation of the network predictions. After training, the analysis of a single plan image took 7.6s corresponding to around 9 min for a full 3D stack. The AI network identified bounding boxes around individual events in each plan independently. We then post-processed the stack using a MATLAB code to link each detected area between z-plan. 3D connected components labelling were used for identifying the same apoptosis/mitosis event. In order to tolerant the slight differences in the xy position of the same event in the consecutive z slices, a 2µm x 2µm bounding box surrounding each xy centre point was used for 3D connected components labelling. Prior to the labelling, we applied median filtering [1×1×3] in z direction to reduce false detections. Finally, we computed the coordinates of the centre point of each 3D box. We plotted the localization map using sphere centred on each of the event coordinates. We measured the confusion matrix by comparing the overlay of the predicted bounding box to the ground truth (**Supplementary Figure 6a**).

### Deep Learning-based classification of organoid shapes

The raw images of global organization of organoids based exclusively on the actin (Phalloidin) and nuclei (DAPI) staining were analyzed using deep learning algorithms developed for image classification. Manually, we distinguished 2 classes of organoids: ‘O’ shaped organoids displaying a spherical global shapes and ‘B’ shapes for which an external streak can be clearly distinguished surrounding by 2 cellular lobes tending to join. We analyzed 132 (2048*2048*71 planes) 3D images. We used DenseNet121^5^, a Convolutional Neural Network (CNN) algorithm for classify the 3D images into two classes: B-shape and O-shape (see Supplemental Figure 6B) on a Geforce RTX 2080 Ti NVIDIA GPU with 11Go of graphical memory. To accommodate the memory constraint of the graphics card, we arbitrarily resized the whole dataset to 128*128*64, a size where the structures are still roughly distinguishable. A 5-fold cross-validation was performed with a 60/20/20 split for training, validation and test set definition. Each fold was launched with a DenseNet121^5^ implementation with Adam optimizer^6^. To counterbalance large imbalance between the two classes, each training iteration was launched with a batch consisting of as many ‘O’ shapes as ‘B’ shapes. Training procedures were performed over 50 epochs, and the weights resulting in a maximum of balanced accuracy score on the validation set are saved and used for the final test. After training, the Predicting the classes among a set of 130 took less than 1s. By gathering predictions on each fold, we obtained 99% of accuracy score. Subsequently, each ‘O’ shape organoids were submitted to AI-based segmentation of internal streaks.

### Deep learning-based segmentation of internal streaks

Classified ‘O’ organoids were proceeded for the segmentation of internal streak based only on morphological staining (DAPI and Phalloidin) to permit detection of morphological patterns independently of genes expression level (**Supplementary Figure 6c**). The 3D dataset was first resized to 1024 * 1024 * 71, and each image plane was manually segmented using 3D Slicer software^7^, giving a ground truth mask. Original image and corresponding masks were again resized to 256*256*71 for faster testing of AI algorithms. The same 5-fold cross-validation procedure described previously were applied using 3D U-net^8^ convolutional network. For training, 128*128*64 tiles of images were picked for batch generation, with a balancing operation between tiles with segmented streak and tiles without segmented streak. Inference process produces probability maps of size 256 * 256 * 71 for each image. An arbitrary threshold of 0.5 was applied to each map, resulting in a binary streak segmentation mask. Once the network trained, the streak detection with the generation of the corresponding mask take approximatively 5sec for each organoid. A 3D connected components algorithm was used to label each voxel of the binary masks. For feature extraction, another rescaling operation was performed in order to match the original image size (for x,y dimensions) and a isotropic voxel size (for z dimension). Most of segmented objects seem relevant for internal streaks as showed in Figure 6 and Supplemental figure 6. Supplementary Table 2 describes the list of the morpho-descriptors that we extracted from the segmented streaks.

## REFERENCES

1. Kim, J., Koo, B. K. & Knoblich, J. A. Human organoids: model systems for human biology and medicine. Nature Reviews Molecular Cell Biology (2020) doi:10.1038/s41580-020-0259-3.

2. Takebe, T. & Wells, J. M. Organoids by design. Science (80-.). 364, 956–959 (2019).

3. Kratochvil, M. J. et al. Engineered materials for organoid systems. Nat. Rev. Mater. 4, 606–622 (2019).

4. Rossi, G., Manfrin, A. & Lutolf, M. P. Progress and potential in organoid research. Nat. Rev. Genet. 19, 671–687 (2018).

5. O’Connell, L. & Winter, D. C. Organoids: Past learning and future directions. Stem Cells Dev. 29, 281–289 (2020).

6. Vives, J. & Batlle-Morera, L. The challenge of developing human 3D organoids into medicines. Stem Cell Research and Therapy (2020) doi:10.1186/s13287-020-1586-1.

7. Busslinger, G. A. et al. The potential and challenges of patient-derived organoids in guiding the multimodality treatment of upper gastrointestinal malignancies. Open Biol. (2020) doi:10.1098/rsob.190274.

8. Lukonin, I. et al. Phenotypic landscape of intestinal organoid regeneration. Nature (2020) doi:10.1038/s41586-020-2776-9.

9. https://bioengineeringcommunity.nature.com/posts/intestinal-regeneration-lessons-from-organoids.

10. Rios, A. C. & Clevers, H. Imaging organoids: A bright future ahead. Nature Methods (2018) doi:10.1038/nmeth.4537.

11. Dekkers, J. F. et al. High-resolution 3D imaging of fixed and cleared organoids. Nat. Protoc. 14, 1756–1771 (2019).

12. Eismann, B. et al. Automated 3D light-sheet screening with high spatiotemporal resolution reveals mitotic phenotypes. J. Cell Sci. 133, 1–20 (2020).

13. Voleti, V. et al. Real-time volumetric microscopy of in vivo dynamics and large-scale samples with SCAPE 2.0. Nat. Methods (2019) doi:10.1038/s41592-019-0579-4.

14. Yang, B. et al. Epi-illumination SPIM for volumetric imaging with high spatial-temporal resolution. Nat. Methods (2019) doi:10.1038/s41592-019-0401-3.

15. Galland, R. et al. 3D high-and super-resolution imaging using single-objective SPIM. Nat. Methods 12, 641–644 (2015).

16. Lancaster, M. A. et al. Cerebral organoids model human brain development and microcephaly. Nature (2013) doi:10.1038/nature12517.

17. Karzbrun, E., Kshirsagar, A., Cohen, S. R., Hanna, J. H. & Reiner, O. Human brain organoids on a chip reveal the physics of folding. Nat. Phys. (2018) doi:10.1038/s41567-018-0046-7.

18. Hříbková, H., Grabiec, M., Klemová, D., Slaninová, I. & Sun, Y. M. Calcium signaling mediates five types of cell morphological changes to form neural rosettes. J. Cell Sci. 131, (2018).

19. Fedorova, V. et al. Differentiation of neural rosettes from human pluripotent stem cells in vitro is sequentially regulated on a molecular level and accomplished by the mechanism reminiscent of secondary neurulation. Stem Cell Res. (2019) doi:10.1016/j.scr.2019.101563.

20. Meinhardt, A. et al. 3D reconstitution of the patterned neural tube from embryonic stem cells. Stem Cell Reports (2014) doi:10.1016/j.stemcr.2014.09.020.

21. Chandrasekaran, A. et al. Comparison of 2D and 3D neural induction methods for the generation of neural progenitor cells from human induced pluripotent stem cells. Stem Cell Res. (2017) doi:10.1016/j.scr.2017.10.010.

22. Jones, T. R. et al. CellProfiler Analyst: data exploration and analysis software for complex image-based screens. BMC Bioinformatics 9, 482 (2008).

23. Huang, G., Liu, Z., Van Der Maaten, L. & Weinberger, K. Q. Densely connected convolutional networks. in Proceedings - 30th IEEE Conference on Computer Vision and Pattern Recognition, CVPR 2017 (2017). doi:10.1109/CVPR.2017.243.

24. Driehuis, E. et al. Pancreatic cancer organoids recapitulate disease and allow personalized drug screening. Proc. Natl. Acad. Sci. U. S. A. (2019) doi:10.1073/pnas.1911273116.

25. Yoshida, S., Miwa, H., Kawachi, T., Kume, S. & Takahashi, K. Generation of intestinal organoids derived from human pluripotent stem cells for drug testing. Sci. Rep. (2020) doi:10.1038/s41598-020-63151-z.

26. Beccari, L. et al. Multi-axial self-organization properties of mouse embryonic stem cells into gastruloids. Nature (2018) doi:10.1038/s41586-018-0578-0.

27. Sorrentino, G. et al. Mechano-modulatory synthetic niches for liver organoid derivation. Nat. Commun. (2020) doi:10.1038/s41467-020-17161-0.

28. Kumar, S. V. et al. Kidney micro-organoids in suspension culture as a scalable source of human pluripotent stem cell-derived kidney cells. Dev. (2019) doi:10.1242/dev.172361.

29. Perche, F. & Torchilin, V. P. Cancer cell spheroids as a model to evaluate chemotherapy protocols. Cancer Biol. Ther. (2012) doi:10.4161/cbt.21353.

## Bibliography

1. Galland, R. et al. 3D high- and super-resolution imaging using single-objective SPIM. Nat. Methods 12, 641–644 (2015).

2. Seglen, P. O. Preparation of Isolated Rat Liver Cells. Methods Cell Biol. (1976) doi:10.1016/S0091-679X(08)61797-5.

3. Niblack, W. An introduction to digital image processing. An Introd. to Digit. image Process. (1986) doi:10.1201/9781315123905-1.

4. Legland, D., Arganda-Carreras, I. & Andrey, P. MorphoLibJ: Integrated library and plugins for mathematical morphology with ImageJ. Bioinformatics (2016) doi:10.1093/bioinformatics/btw413.

5. Huang, G., Liu, Z., Van Der Maaten, L. & Weinberger, K. Q. Densely connected convolutional networks. in Proceedings - 30th IEEE Conference on Computer Vision and Pattern Recognition, CVPR 2017 (2017). doi:10.1109/CVPR.2017.243.

6. Kingma, D. P. & Ba, J. L. Adam: A method for stochastic optimization. in 3rd International Conference on Learning Representations, ICLR 2015 - Conference Track Proceedings (2015).

7. Fedorov, A. et al. 3D Slicer as an image computing platform for the Quantitative Imaging Network. Magn. Reson. Imaging (2012) doi:10.1016/j.mri.2012.05.001.

8. Ronneberger, O., Fischer, P. & Brox, T. U-Net: Convolutional Networks for Biomedical Image Segmentation [2015; First paper exploring U-Net architecture.]. in International Conference on Medical image computing and computer-assisted intervention (2015).

