## Supplementary info for "High content 3D imaging method for quantitative characterization of organoid development and phenotype"

### Supplementary Materials

#### A- Supplementary Figures

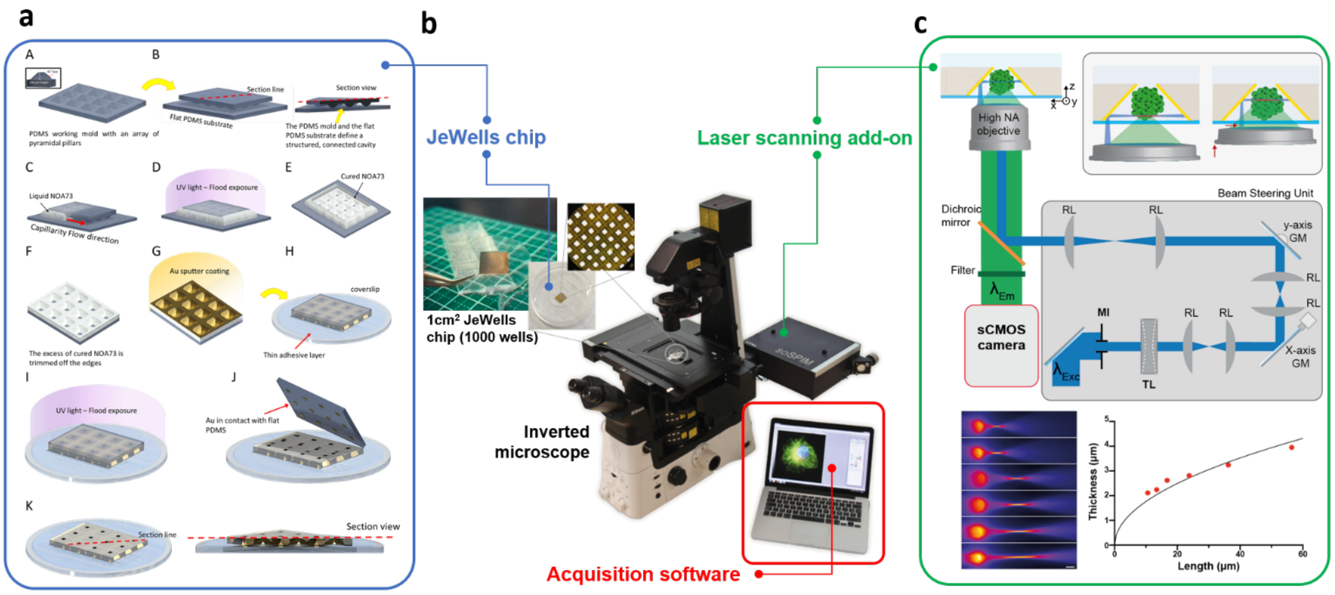


**Supplementary Figure 1: Overview of the platform. a.**Schematic representation of the fabrication steps of the Jewells chips. A) Scheme of the PDMS mold, the inset on the top left corner shows a detail of the geometry of the truncated pyramids. B) The PDMS working mold is flipped and placed on top of a flat cut of PDMS, the red dashed line shows a sectioning (right side of the figure) of the assembly where the cavity comprised between the two PDMS parts is indicated. C) UV curable resin (NOA73 in this case) is poured next to one edge and by capillarity action fills the cavity (the red arrow shows the direction of flow). D) After complete filling of the cavity, NOA73 is cured by UV flood exposure. E) The PDMS mold is peeled-off, leaving the cured NOA73 on the flat PDMS cut, which serves as a handling substrate. F) Excess of cured NOA73 from the 4 edges is trimmed away. G) Au thin film coating (by sputter-coating). H) A standard coverslip is coated with thin layer of NOA73, which is then partially cured to become solid while retaining adhesive properties; the Au-coated film is flipped and pressed on the pre-cured NOA73. I) UV flood-exposure (from the coverslip side) to fully cured the thin adhesive NOA73 layer. J) Peeling-off of the flat PDMS handling substrate, which removes the Au film in contact and leaves the JeWell structured film adherent to the coverslip and with the top of the pyramidal micro-wells open. K) Final JeWell as a reflective structured NOA73 film on a glass coverslip with open access at the top; the red dashed line shows a sectioning of the JeWell pyramidal reflective microwells (right side of the picture). **b**. Image of a functional soSPIM set-up composed of a standard inverted microscope, the soSPIM beam steering unit plugged at the back port of the microscope body (green box.), the micro-fabricated device positioned on the microscope stage (blue box), and the soSPIM software (red box) which enables to control and synchronize the optical elements to perform 3D imaging. **c.** soSPIM optical setup and acquisition principles. Schematic representation of the soSPIM principle showing a sample holder comprising 45° micro-mirrored cavities deposited on a glass coverslip and an excitation beam–steering unit mounted on a conventional inverted microscope. Beam steering[1] is performed by reflection on two galvanometric mirrors (GM) and a focal tunable lens (TL) all conjugated by relay lenses (RL) to the objective back focal plane. A motorized iris (MI) allows adjusting the laser beam diameter to control the light -sheet parameters (thickness and length). Upper right: cartoon illustrating the 3D acquisition principle of the soSPIM system. The light-sheet position onto the 45° mirror is synchronously adjusted with the objective axial position and the tunable lens to maintain the excitation and detection plane superposed and the light-sheet thinnest part at the sample position for 3D imaging. Light-sheet thickness (FWHM) as function of its length (FWHM) obtained for different iris opening within the beam steering unit (red dots). The light sheet parameters have been computed from the images of the light-sheet after reflection onto a 45° mirror into a well filled by a fluorescent solution represented. The light-sheet thicknesses correspond to the minimum width of the beams perpendicular to their propagation direction as fitted by a gaussian function, and the light sheet length to the width of the beams along their propagation direction as fitted by a gaussian beam. The black line represents the theoretical relationship in between the thickness and the length a focalized gaussian beam at 488 nm.

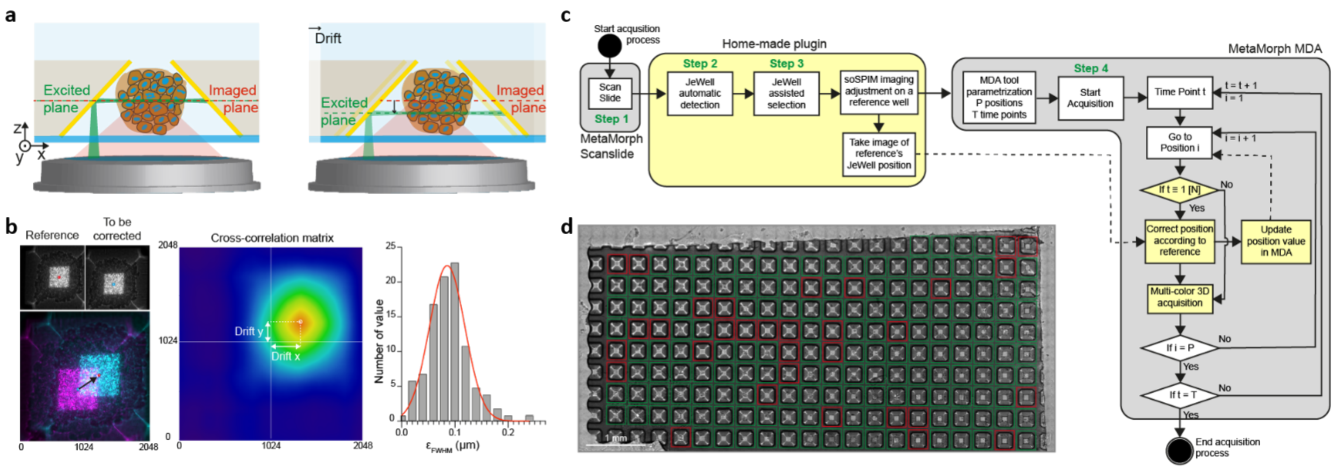


**Supplementary Figure 2:**

**JeWells re-positioning tool and Acquisition workflow. a.** Schematic representation of the relation between the XY mirror position and the light-sheet illumination depth created by a laser beam reflected on the 45° mirror. A spatial drift of the device perpendicular to the 45° mirror axis (x-axis on the schema) leads to an equivalent displacement of the light-sheet along the z-axis, whereas the imaged plane remains at the same position, corrupting the image quality. Lateral drift needs to be corrected to ensure optimal 3D imaging of long-term and HCS-like multi-position acquisitions. **b.** Principle of the cross-correlation-based repositioning system. Left: Comparison between a reference position and the drifted position to correct, and the overlay before (Magenta) and after (Cyan) correction. Middle: Cross-correlation matrix of the two brightfield images represented on the left panel. Right: Histogram of the light-sheet width broadening after the cross-correlation-based repositioning obtained from 100 random displacements (in a range of +/- 3 µm) perpendicular to the mirror axis (x-axis in a). It resulted to a mean broadening of 0.085 +/- 0.033 µm (<3% of a 2.5 µm thick light-sheet). **c.** FlowChart representation of a high-content imaging acquisition process: A bright-field preview of the full chip containing JeWell array is performed using the scanslide module of MetaMorph (Step 1). The automatic detection of the JeWells to image (Step 2) and their interactive selections (Step 3) is handled through a home-made plugin integrated into MetaMorph environment with the soSPIM beam steering unit control (yellow box). The acquisition process is then steered using the Multi-Dimensional Acquisition module of MetaMorph (Step 4) in which journals calling for home-made routines are integrated enabling Jewells accurate re-positioning and soSPIM 3D imaging (yellow rectangles). The acquisition process ends once all the positions have been acquired at every time-points. **d.** Left: ScanSlide of a soSPIM device containing 299 JeWells with their automatic detection (ROI) and assisted selection (green/red). The overall ScanSlide acquisition, detection and selection processes took 25 min.

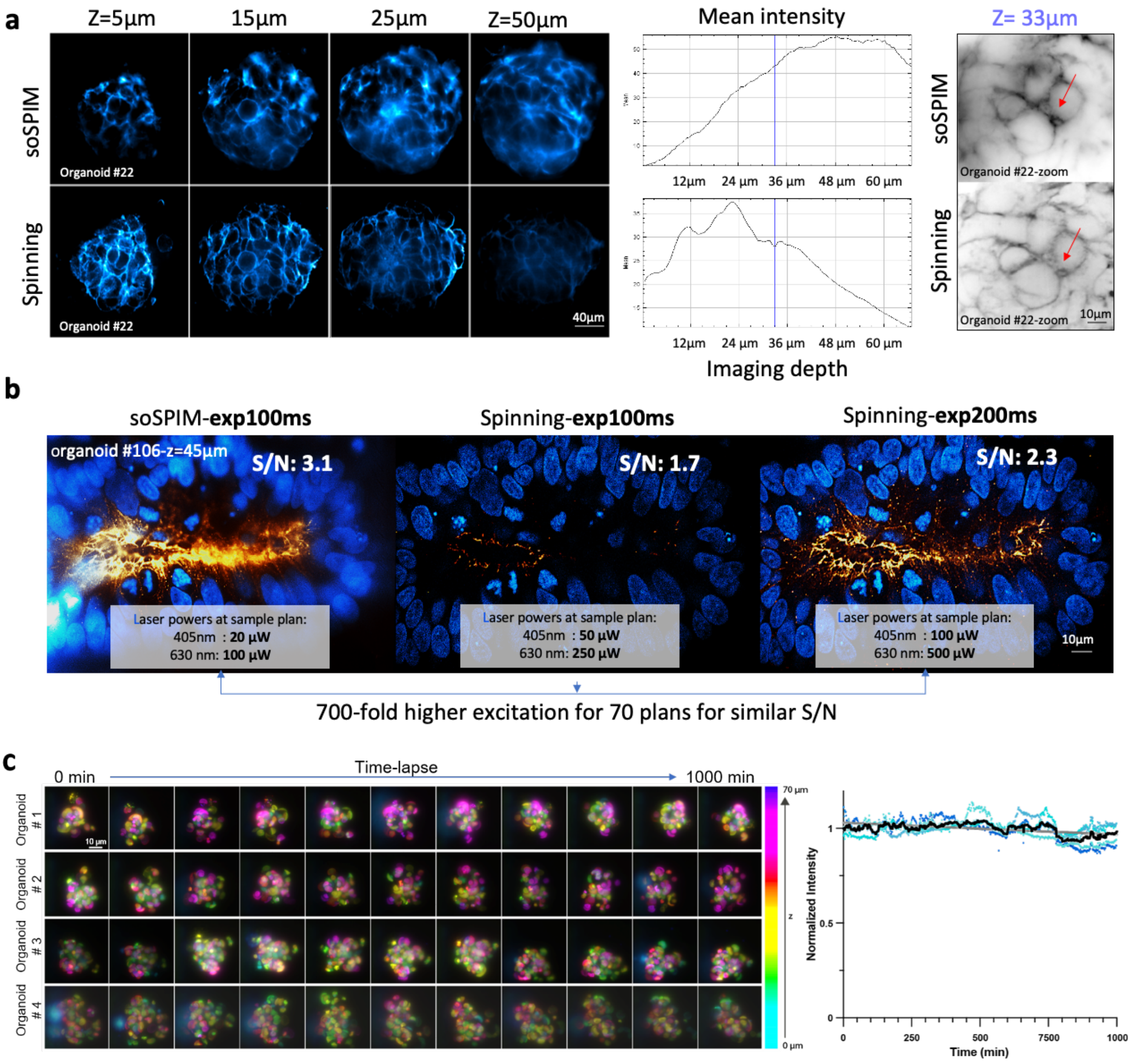


**Supplementary Figure 3:**

**Comparison of soSPIM with spinning-disc microscopy (Yokogawa W1) and soSPIM bleaching measurement. a.** We used the same microscope body (Nikon Eclipse), camera (Hamamatsu Orca Flash 4), laser launch (Oxxius laser diodes 405, 488, 630 nm) and objective (60x WI 1.2NA) to compare the imaging quality of the soSPIM approach with the spinning-disc. Out of focus signal rejection was superior for spinning-disc between 0 and 25 µm above the coverslip. Deeper than 25 µm, the light-sheet illumination superseded the spinning-disc, collecting more signal. This is an expected characteristic when comparing confocal to light sheet microscopy. **b.** Comparison of the total laser power received by an organoid during the acquisition of 70 z-Stacks. For equivalent signal to noise ratio, the sample the light exposure is 700 time less for soSPIM than for spinning. The laser power is measured at the focal plane of the objective. **c.** Left : depth color-coded maximum intensity projection of selected time-points of a time-lapse soSPIM acquisition performed on 4 different HEP3B spheroids stably expressing H2B-GFP. 3D stacks of 70 planes (every 1 µm) were acquired every 90 seconds for 721 time-points corresponding to a whole acquisition duration of 18 h (1080 min). Right : normalized mean intensity of the spheroids signal over the whole acquisition duration (dots) and mean curves (black line) fitted with a single exponential decay (grey line) illustrating the low to no bleaching of the signal along the time-course of the acquisition (half-life = 11078 +/- 900 min). The difference of the mean intensity on a volume encompassing the spheroid and a volume outside of the spheroid were computed for each time-points and positions. The signal was then normalized according to its mean value for all the time-points. All these effects explain why we could perform prolonged live imaging in 3D using the acquisition platform.

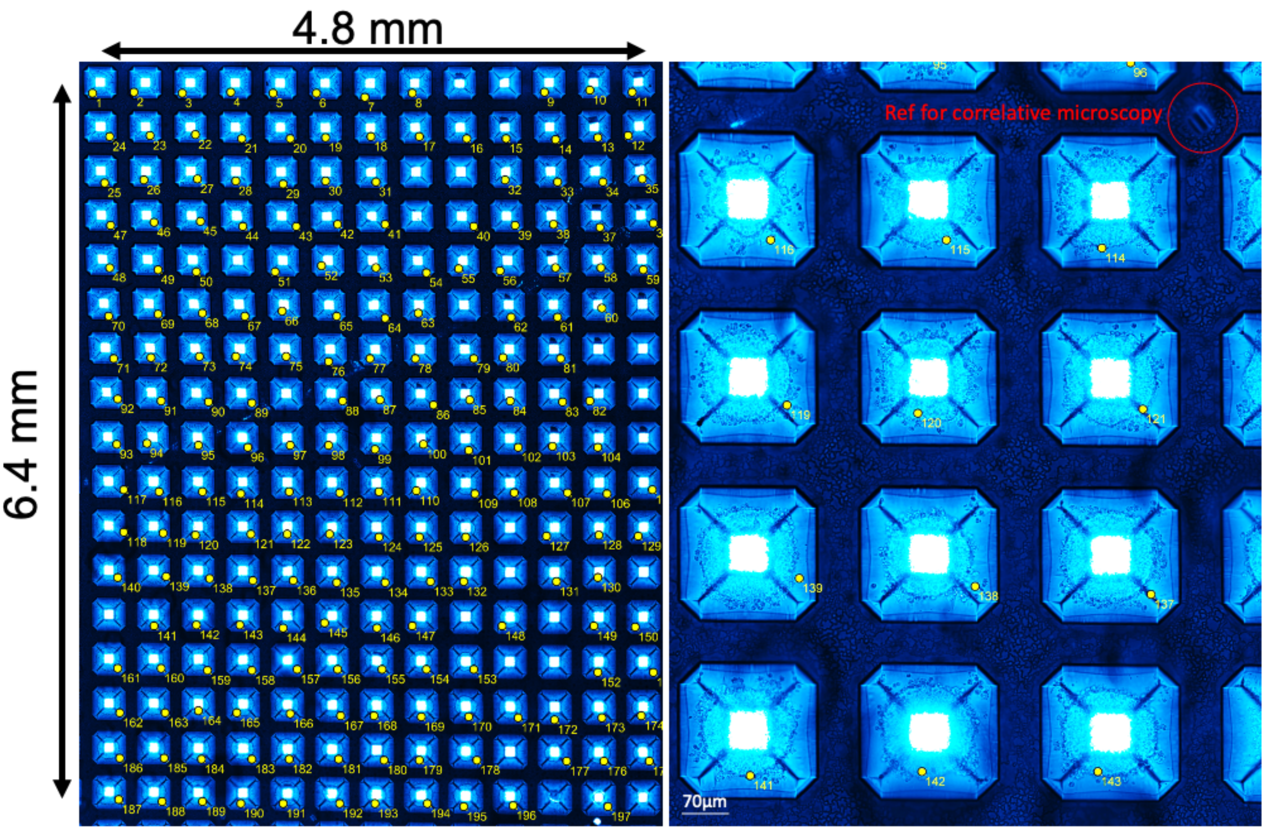


197/221 = 90% of occupancy. 1 organoid per JeWell

**Supplementary Figure 4:**

**Scanned view of the Jewell chip with 90% Jewell single organoid occupancy. a.** Large scale view of the 221 Jewells on a chip under a phase contrast microscope (20x). The Jewells appear as squares with 4 slanted sides. The numbers reference the Jewells that are populated by a single organoid. Organoids in each Jewell, obtained after 8 days of differentiation process of hESC to neuroectoderm, appear as a ball surrounded by cellular debris. 90% of the Jewells displayed an organoid. Reference marks were made by onsite laser engraving, using the soSPIM excitation hardware. It permits the repositioning of the plate on different acquisition systems or after any manipulation of the plate outside the platform (medium exchange, ECM addition…). A specific ID is attributed to each organoid for correlative microscopy.

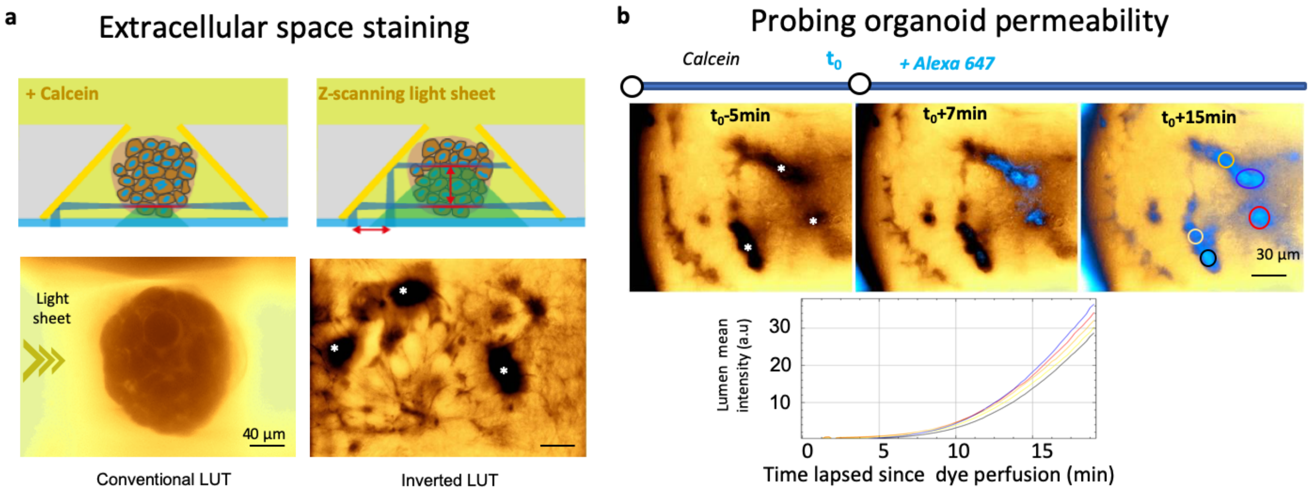


**Supplementary Figure 5:**

**3D live imaging using extracellular dyes. a.** Schematic of the 3D imaging scheme in a Jewell. Synchronizing the position of the imaging plane (Z position of the objective) with the position of the excitation light-sheet (position of the laser on the mirrors) permits the automated 3D imaging of the organoids in the Jewells. Introducing calcein as an extracellular dye, allows revealing the intercellular spaces and luminal cavities (marked by *) in the 3D culture (inverted LUT). The illumination comes from one side (left here). **b**. We introduced a second no-permeant dye (Alexa 647) and used high speed imaging (12 stacks/min) to monitor the dynamics of infiltration of the dye in the 3D culture.

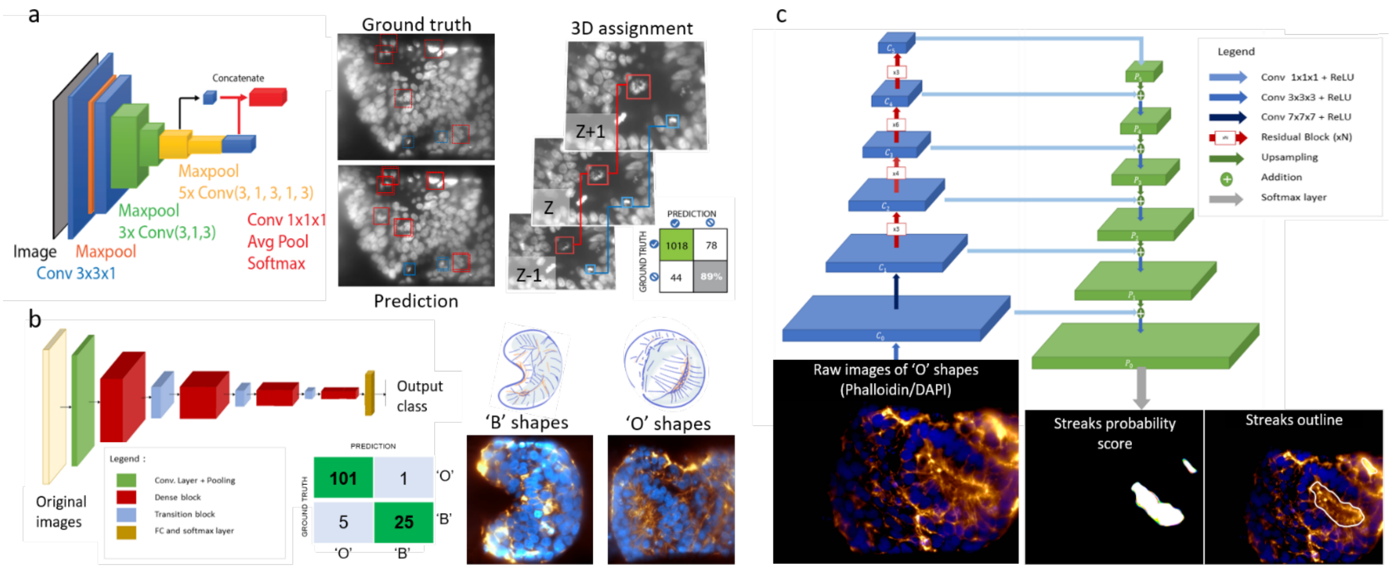


**Supplementary Figure 6: AI based analysis used in the workflow. a.** Schematic description of the AI based analysis pipeline for mitosis and apoptosis detection using the neural network YOLOv2 published by ZeroCostDL4Mic with default settings using our own GPU resources (NVIDIA Quadro RTX6000 24GB). The ground truth has consisted of a training dataset of mitosis and apoptosis manually pinpointing by drawing bounding boxes around the central plane of the apoptosis (Blue) and of the mitosis (Red). A total of 344 mitosis and 1123 apoptosis were manually picked for the training, augmented to a final training set of 4128 mitosis and 13476 apoptosis (See online methods). We used 90% of this set for the training and 10% for validation of the network. The AI detection results (Prediction) showed an accuracy of 89% as compared to the ground truth. The prediction was obtained by processing each z slice independently, it has been subsequently post-processed to obtain the 3D assignment of all events (See online methods). **b**. Illustration of the network architecture used for the whole organoid classification task, corresponding to an 3D adaptation of Densenet121 model[2]. It has resulted in a 99% accuracy as compared to ground truth for the classification of organoids in 'B' and 'O' shapes (representative images). c. Illustration of Features pyramid network (FPN) architecture with successive Resblocks in the backbone (C0-C5) of the bottom-up pathway (blue). The remain features maps (P0-P5) was generated by a combination of convolution operation applied to the feature maps of the bottom up pathway and upsampling transformation applied on each level of the pyramid (green). The probability score and segmentation output (Streaks outline) was given by applying softmax layer on P0.

#### B- Supplementary Movies:

**Movie 1:**

**Live imaging using calcein in extracellular medium**. Organoid made from hIPSC in differentiation process to hepatoblast (day10 in Jewells), counterstained with calcein (inverted LUT). One 3D z-stack every 15 min (only the median plan of the organoid is displayed).

**Movie 2:**

**Parallelized live soSPIM acquisition of differentiating organoids.** Organoids made from hESC-lifeActGFP-H2BmCherry in differentiation process to neuroectoderm in Jewells. Green : lifeAct, Purple : histone-H2B. 15 different organoids every 15 min. Top line for high Z, middle line for median Z, bottom line for low Z.

**Movie 3:**

**Eight days time-lapse recording of Neuroectoderm differentiation.** Organoid made from hESC-lifeActGFP in differentiation process to neuroectoderm (from day 2 to day 8 in Jewells) (inverted gray LUT) (median plan of the organoid). Yellow line is corresponded to a rosette-shape organization around an internal streak.

**Movie 4:**

**Multi light sheet mode of the soSPIM of one organoid labelled with phalloidin (actin).** Three light sheets are created successively on the left, bottom, and right mirror (symbolled by lines). The three images can be superimposed to obtain a best contrast image (median plan of the organoid). This imaging mode has not been implemented automatically.

**Movie 5:**

**3D stacks of one organoid of the screening pool and multi light sheet mode.** Organoid made from hESC in differentiation process to neuroectoderm (day8 in Jewells). Thanks to the identified positioning of each organoid, it is possible to re-acquire some of organoids of interest, i.e with the multilight sheet mode. Blue : DAPI, Magenta : Sox2, Gold : Phalloidin.

**Movie 6:**

**AI probability and segmentation of streaks in an organoid.** Organoid made from hESC in differentiation process to neuroectoderm (fixation and staining at day 8 in Jewells). 3d stack images were acquired in soSPIM mode for Actin (Phalloidin, gold) and nuclei (DAPI, blue). Left: AI probabilities map for each pixel to belong to a streak, based on the actin staining. Right: segmentation over raw images.

**Movie 7:**

**AI based segmentation of a streak in 3D.** The streak is the region encompassed within the white volume that results from the thresholding of the AI scores showed in Movie 7. Note that the detected region corresponds to the core of the rosette and does not encompass the surrounding cells. The segmentation was performed on the actin staining images, but it corresponds to the N-cadherin positive region of the rosettes.

**Movie 8:**

**96 primary rat hepatocytes spheroids and z stack .** Primary rat hepatocytes were seeded in the jewell and left for aggregation during 2 days. We then fixed them and stained for actin (Phalloidin, grey) and MRP2 (Gold). MRP2 is an apical transporter that is recruited at the apical poles of the hepatocytes. It is a reporter for the bile canaliculi. The medial Z plane is displayed for each of the 96 spheroids on the left, and a z-stack of one of the 96 spheroids on the right.

#### C- Bibliography :

1. Galland, R. *et al.* 3D high- and super-resolution imaging using single-objective SPIM. *Nat. Methods* **12**, 641–644 (2015).

2. Huang, G., Liu, Z., Van Der Maaten, L. & Weinberger, K. Q. Densely connected convolutional networks. in *Proceedings - 30th IEEE Conference on Computer Vision and Pattern Recognition, CVPR 2017* (2017). doi:10.1109/CVPR.2017.243.

| Primers | Forward | Reverse |
|---|---|---|
| Human SIP1 | CAAGAGGCGCAAACAAGCC | GGTTGGCAATACCGTCATCC |
| Human OTX2 | CACTTCGGGTATGGACTTGC | CGGGTCTTGGCAAACAGTG |
| Human PAX2 | TGTCAGCAAAATCCTGGGCAG | GTCGGGTTCTGTCGTTTGTATT |
| Human PAX5 | ACCAGCAGGACAGGACATGG | TCCACTATCCTCTGGCGGAC |
| Human FGF8 | AAAGCTCATCGTGGAGACGG | GCCCTCGTACTTGGCATTCT |
| Human IRX3 | CTCTCCCTGGTGGCGTTCTT | GTGTCCCTTCCTTCTCCATGTG |
| *SOX2* | GAACCAGCGCATGGACAGTTAC | TGTAGGTCTGCGAGCTGGTCAT |
| *PAX6* | TCTTTGCTTGGGAAATCCG | CTGCCCGTTCAACATCCTTAG |
| *N-CADHERIN* | CCACCTTAAAATCTGCAGGC | GTGCATGAAGGACAGCCTCT |
| *E-CADHERIN* | CGAGAGCTACACGTTCACGG | GGGTGTCGAGGGAAAAATAGG |
| *NANOG* | CAAAGCTTGCCTTGCTTTGAAGC | AAGGAAGAGGAGAGACAGTCTCC |
| *GAPDH* | ATGCTGGCGCTGAGTACGTCGTG | GTGCTAAGCAGTTGGTGGTGCAG |

**Table 1: List of primers sequence used for RT qPCR.**

| Level | Feature | Units |
|---|---|---|
| **Whole organoid level** | Diameter | µm |
| | Morphoclass | number |
| | Number of cells | count |
| | Mitosis index | Percentage vs. number of cells |
| | Apoptosis index | Percentage vs. number of cells |
| **Multicellular level (i.e streak)** | Volume | $\mu m^3$ |
| | Surface area | $\mu m^2$ |
| | Length of skeleton | µm |
| | 3D positioning (center) | µm |
| | Sphericity | a.u |
| | Thickness | µm |
| | Shapes descriptors… | various |
| **Single cells level** | Volume of nucleus | $\mu m^3$ |
| | Surface area of nucleus | $\mu m^2$ |
| | Length of nucleus skeleton | µm |
| | 3D positioning (center) | µm |
| | Sphericity | a.u |
| | Shapes descriptors… | various |
| | Texture descriptors… | various |
| | Expression levels of different markers | a.u |

**Table 2: Summary of the morphological features extracted with the workflow and fed to the database**. Each *shape and texture descriptors tens of different parameters not detailed here. This list is not exhaustive and could be completed regarding the biological context.*
